# A Comprehensive Conceptual and Computational Dynamics Framework for Autonomous Regeneration of Form and Function in Biological Organisms

**DOI:** 10.1101/2022.09.05.506675

**Authors:** Sandhya Samarasinghe, Tran Nguyen Minh-Thai

## Abstract

In biology, regeneration is a mysterious phenomenon that has inspired self-repairing systems, robots, and biobots. It is a collective computational process whereby cells communicate to achieve an anatomical set point and restore original function in regenerated tissue or the whole organism. Despite decades of research, the mechanisms involved in this process are still poorly understood. Likewise, the current algorithms are insufficient to overcome this knowledge barrier and enable advances in regenerative medicine, synthetic biology, and living machines/biobots. We propose a comprehensive conceptual framework for the engine of regeneration with hypotheses for the mechanisms and algorithms of stem cell-mediated regeneration that enables a system like the planarian flatworm to fully restore anatomical (form) and bioelectric (function) homeostasis from any small- or large-scale damage. The framework extends the available regeneration knowledge with novel hypotheses to propose collective intelligent self-repair machines, with multi-level feedback neural control systems, driven by somatic and stem cells. We computationally implemented the framework to demonstrate the robust recovery of both anatomical and bioelectric homeostasis in an *in silico* worm that, in a simple way, resembles the planarian. In the absence of complete regeneration knowledge, the framework contributes to understanding and generating hypotheses for stem cell mediated form and function regeneration which may help advance regenerative medicine and synthetic biology. Further, as our framework is a bio-inspired and bio-computing self-repair machine, it may be useful for building self-repair robots/biobots and artificial self-repair systems.

**Summary:** A conceptual framework for the machinery of self-repair in living systems that enables a synthetic organism to accurately regenerate form and function from any disturbance and damage.

## INTRODUCTION

In biology, regeneration refers primarily to morphological processes that characterise the plasticity of the phenotype of traits that allow multi-cellular organisms to repair themselves (self-repair) and maintain the integrity of their morphology (form or anatomical pattern), and their physiological state (function). This is a process of collective action, through cellular computing in living systems, in the form of soft robotics, that displays a high-level of complex adaptive decision-making. With body-wide immortality, Hydra (tiny aquatic animals) and planarians (free-living flatworms) are models of adaptive regeneration [1]. After being injured, cells in any amputated fraction fully restore tissues and organs to their previous anatomical and physiological state. The regeneration of organs is widespread in animals such as snails, axolotls (an amphibian known as a ‘walking fish’), and zebrafish [2]. In a related context, some animals can reproduce asexually through fragmentation (e.g., starfish, some worms, fungi, plants, and lichens), budding (e.g., yeast, Hydra) or fission (e.g., Bacteria, Protists, Unicellular Fungi) [3]. For example, a planarian mother will narrow and split in the middle. Each half creates a new head or tail to form two copies of the original. Despite decades of research on regeneration, how animals reproduce and maintain their morphology and physiological state (e.g., bioelectric homeostasis) remains largely unknown. Living machines and soft robotics require testable models to explain the methods and mechanisms used to maintain the correct morphological and physiological state of the organism [4]. This lack of knowledge has impeded the progress of regenerative medicine. This study proposes a conceptual framework for whole organism morphological and physiological regeneration of a simple synthetic worm modelled on planarian regeneration.

A 6 mm long planarian (Fig. 1A) has approximately 0.6 million cells (Fig. 1B) [5], with adult stem cells (shown in yellow) comprising approximately 20 to 30% of all cells distributed across the planarian’s body [6]. Stem cells produce tissue (somatic) cells that, in the process of development of the organism, divide. After incurring damage, stem cells migrate to the location of damage and reproduce all necessary types of stem and somatic (tissue) cells to regenerate lost tissue and organs, completely restoring the organism. How stem cells (in collaboration with the remaining cells) achieve this remarkable recovery is unknown. In a planarian, a severed head, body, or tail takes just two weeks to grow into a fully formed planarian (Fig. 1A). Planaria are also capable of identifying anterior-posterior (A/P) (along the body) and dorsal-ventral (D/V) (across the body) polarity and patterning (form) information via bioelectric signalling [7, 8] meaning that a small piece of planaria can locate the new head and tail correctly as in a normal planarian (Fig. 1A) [9]. How the remaining cells in a worm split into two segments re-establish polarity in the two segments during regeneration is still unclear. Scientists believe that stem and somatic/tissue cells communicate via bioelectricity, but exactly how this occurs is still unclear [10]. A single stem cell, introduced after damage, can reproduce new stem cells that regenerate the whole pattern in an irradiated animal where no stem cells remain [11]. While the body-wide immortality (ability to regenerate from any damage anywhere) of planaria and the process of regeneration in organisms has attracted much research interest, many questions about the mechanisms and algorithms of regeneration remain. Answering these questions may not only lead to new possibilities for human longevity, but also open avenues for designing advanced self-repairing robots and biobots which resemble living systems.

**Fig. 1.**
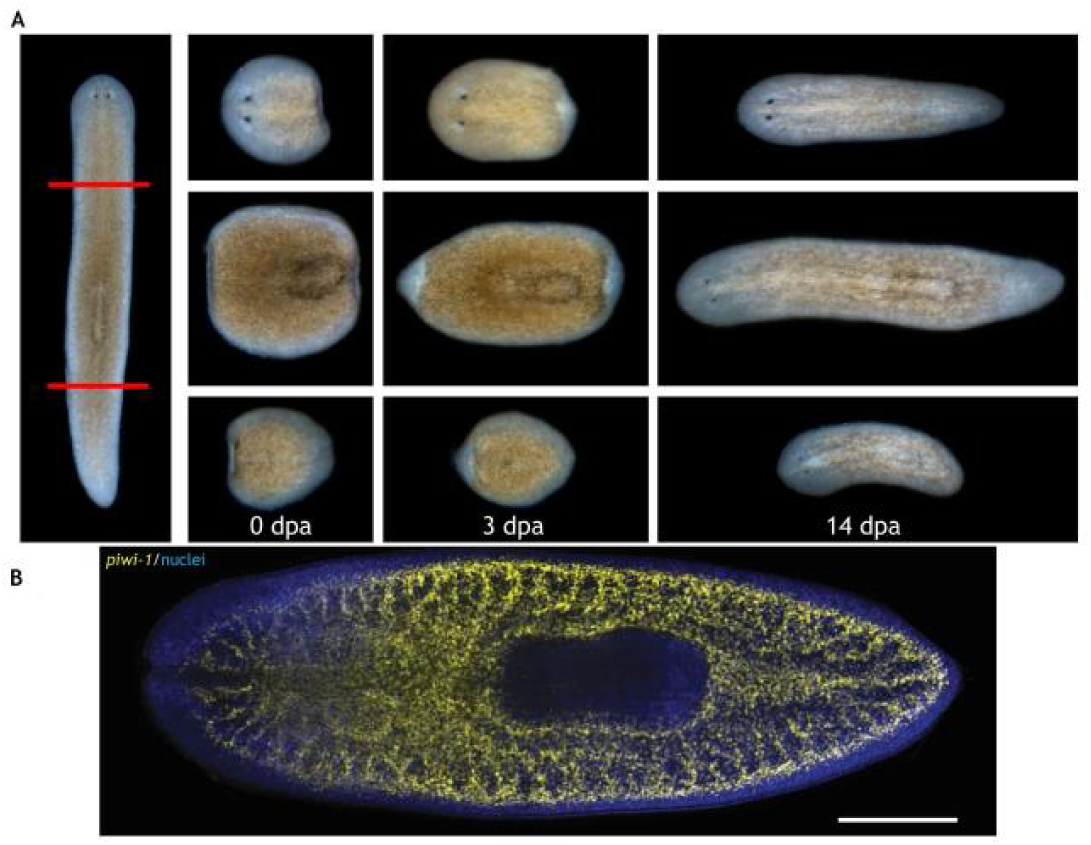
Planarian regeneration and the stem cell system. (A) Regeneration of the head (top), the body (middle), and the tail (bottom), showing the complete regeneration of individual parts into a full worm. Red lines indicate the cutting planes. Days post-amputation (dpa) are observed time points. (B) The distribution of stem cells (yellow); blue indicates somatic (tissue) cells. Scale bar: 500 μm. (Source: Ivankovic et al., 2019, with permission)

Understanding the “mechanisms” and the associated “algorithms” of regeneration are crucial for unravelling how organisms restore the correct anatomy and physiology of organs and the whole organism. In an organism, many physiological functions are sustained by bioelectricity [12]. Upon successful regeneration, it is therefore crucial to maintain the membrane voltage of individual cells across an organism (i.e., body-wide bioelectric equilibrium or homeostasis). However, we still do not understand the mechanisms that coordinate cells into tissue nor how organs are arranged during regeneration. We also do not understand how bioelectricity modulates these responses nor how bioelectric homeostasis is restored by an organism after regeneration. *This study provides a high-level conceptual soft robotic framework for the accurate regeneration of a simple organism. Using stem and somatic cells that communicate via bioelectric signals, the organism is able to restore both anatomical and bioelectric homeostasis*. There are currently no computational models for the accurate maintenance and restoration of morphology (form) or physiology (e.g., bioelectric homeostasis) or both.

Only a few researchers have attempted to create soft-robotics models of biological pattern regeneration using simple synthetic systems. Most have achieved limited success. Furthermore, no one has attempted the restoration of voltage homeostasis from perturbations under normal or damage conditions. Most of the past computational regeneration models have used simple (non-bioelectric) signal interactions: that is, between stem cells and somatic cells or only between stem cells in tissues [4, 13-15]. These models range from simple ones that attempt to restore the total signal received by cells, to agent-based [15] and neural network models [4, 16]. Most of these models require too much information on pattern (form), interactions, computations, and rules for recovering the normal pattern for simple tissue structures. These systems do not fully recover and they do not know when to stop regenerating.

In recent decades, the number of studies on nature-inspired self-repair have increased significantly in robotics [17, 18], software [19-23], and electronics [24, 25]. Most of these studies concentrate on function only repair using information and communication within or without (i.e., internet). A few of these systems can restore form, but to a limited extent (they need new entities or spare parts from ‘outside’). This is a stark departure from biological systems that regenerate new cells from within [6]. However, some of these self-repair models share similarities with the few existing biological regeneration models, the most prominent being short- and long-range communication between entities.

In our previous work, we addressed some of the unresolved issues in biological regeneration [26]. We raised some fundamental questions about regeneration and provided hypothetical answers for regeneration in a simple tissue system. These questions included: How do stem cells sense damage and cooperate with somatic cells to repair local tissue damage? How do stem cells work together in large-scale repair? How do stem cells know what to recover and when to stop? Where could such morphological (pattern) information be stored? How could this information be retrieved during the process of regeneration? Importantly, how does a small body fragment regenerate into a fully formed organism (with correct polarity), to achieve body-wide immortality? In our previous article [26], we provided a simple soft-robotics computational framework to address these issues of regeneration as a collective intelligence problem. That framework, developed for a synthetic (*in silico*) worm consisting of three tissues (a head, body and tail), was based on simple binary communication between cells, indicating their presence or absence, and a simple stem cell arrangement where there was only one stem cell per tissue with a large number of somatic cells. It was also not designed to restore the original function after regeneration.

Our new framework greatly extends the previous one’s capacity by introducing new concepts to address deeper issues of regeneration in biological forms. This framework produces a soft robotic system that dynamically maintains both physiological and morphological homeostasis. It is more biologically realistic with respect to planarian anatomy: it has a greater number of stem cells. This framework also includes cellular communication via bioelectricity which enables it to restore form and function. This new framework raises further questions: How do the stem cells, dispersed throughout a tissue (Fig. 1B), accomplish form regeneration in collaboration with tissue cells? What communication structures are involved in this process? How does a cell collective hold a bioelectric pattern in the form of a body-wide voltage gradient? Finally, how does the organism restore bioelectric homeostasis under normal perturbations due to regular physiological functioning or in regeneration?

To better understand regeneration, our framework incorporates what is known and what is required for accurate regeneration and bioelectric homeostasis that resembles body-wide immortality in planaria. In this new framework, a change in the bioelectric state, due to normal physiological functioning or damage, triggers both appropriate regeneration mechanisms and those that restore bioelectric homeostasis. Our conceptual framework integrates mechanisms and algorithms of pattern and bioelectric homeostasis within a multi-level organisation of bioelectricity mediated computing cell networks (Fig. 2) that form a neural control system that autonomously maintains form and function under any small or large disturbances. We use a simple artificial (*in silico*) worm-like organism to demonstrate the accuracy and robustness of the regeneration framework.

**Fig. 2.**
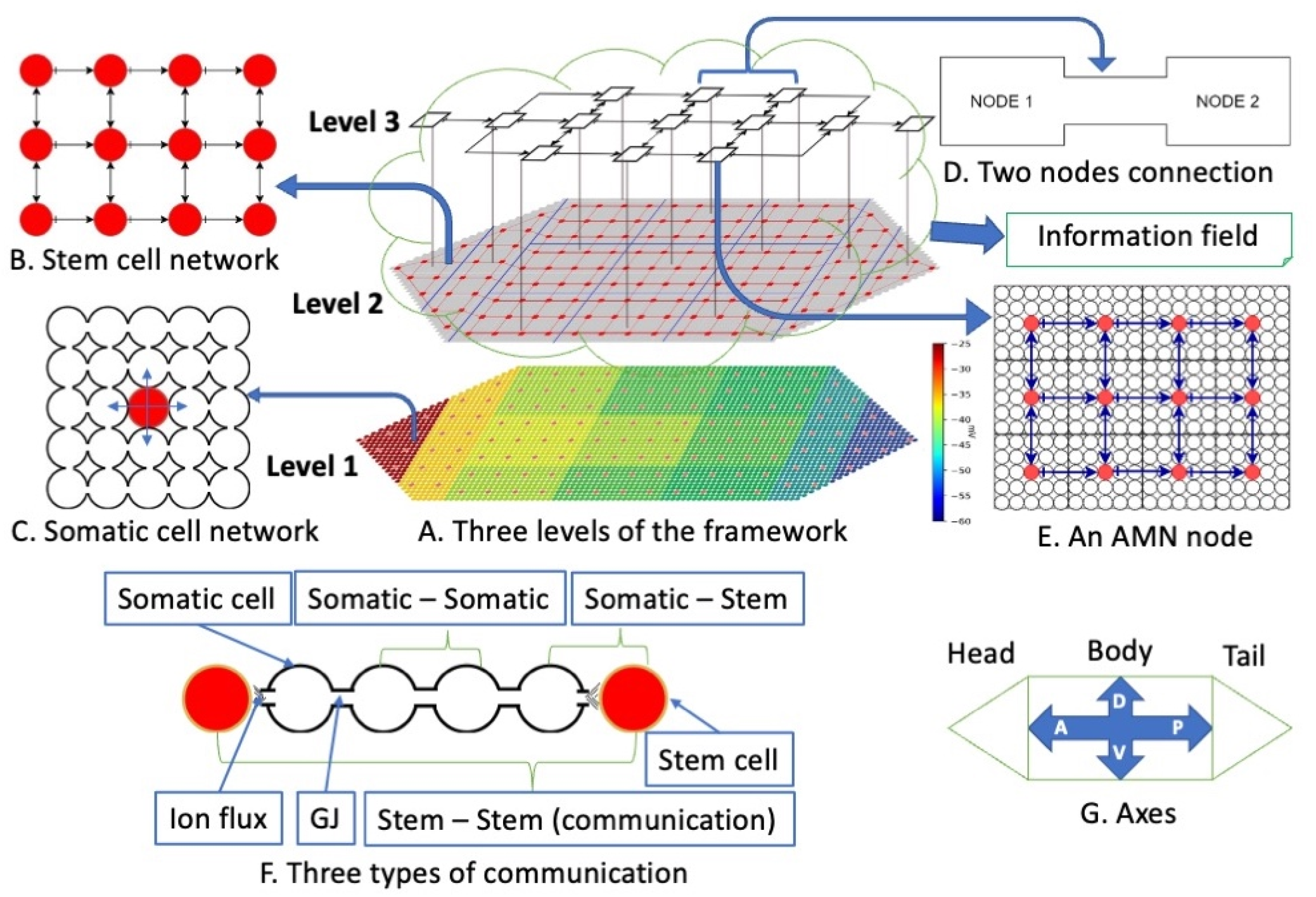
Detailed view of the framework for autonomous restoration of anatomical and bioelectric pattern homeostasis. (A) Three levels of the framework: Level 1- the somatic/tissue cell network (the bottom layer). There are approximately 150 stem cells (red dots) distributed through the organism and 3750 somatic cells surrounding the stem cells (tiny dots filling the whole organism); colours indicate voltage, increasing from green to red as indicated in the scale bar to the right). Level 2 – the stem cell network (the middle) (the engine of regeneration) with an information field that contains minimum pattern and bioelectric information; and Level 3- the global aggregate nodal network (the top). All three levels form computational networks: Levels 1 and 2 are perceptron networks that collectively perform pattern regeneration; Level 3 creates an Associative Memory Network (AMN) (the top) that restores bioelectric homeostasis; all 3 levels cooperate in pattern and bioelectric homeostasis. (B) A detailed view of the stem cell network, showing connections between the stem cells. (C) The somatic cell network in a segment of tissue consisting of a stem cell (red) surrounded by somatic cells. These segments repeat to form the head, body, and tail tissues that constitute the whole pattern. (D) Two nodes of the AMN where the connection represents four Gap Junctions (GJ) connecting four somatic cells in the respective nodes. (E) A detailed view of an AMN node that comprises a segment of a tissue with approximately 300 cells: stem cells (red dots) and somatic cells (white dots). In all networks, arrows indicate the flow of bioelectric communication signals. (F) Three types of communication between cells (red and white cells are stem and somatic cells, respectively): Communication between two somatic cells occurs through GJ; communication between a stem cell and a neighbour somatic cell occurs via ion fluxes; between two neighbour stem cells, communication occurs indirectly, through the GJs of intermediate somatic cells and ion fluxes. (G) The two main axes of the planarian structure: Anterior-Posterior (A/P) and Dorsal-Ventral (D/V).

The proposed soft robotic framework may enable the creation of self-repairing biobots, synthetic living machines, and artificial robots [27-32]. It may also provide a realistic platform for advancing hypotheses to better understand the algorithmic mechanisms of regeneration and promote advancements in regenerative medicine for solving biomedical problems such as congenital disabilities, injury from trauma, cancer, and ageing [33].

Fig. 2 shows the framework, highlighting the concepts of regeneration –the mechanisms and algorithms-using a synthetic worm with three tissues, head, body, and tail (Fig. 2A). It contains novel aspects: First, we assume that the organism maintains a longitudinal bioelectric gradient, from the head to the tail, and a transverse bioelectric gradient, across the body (Fig. 2A Level-1). Second, the organism has a larger number of stem cells (4% of the total) distributed throughout the body (Fig. 2A, Level-2). Third, all communication follows the body-wide bioelectric gradient. Figure 2F provides an example of a local neighbourhood of stem cells consisting of two stem cells (red colour) and four intermediate somatic tissue cells. Tissue cells communicate with each other locally and directly through physical connections called Gap Junctions (GJ). Stem cells communicate with (neighbouring) somatic cells via ion fluxes released to the environment. Accordingly, stem cells communicate with each other indirectly through the GJs of intermediate somatic cells and ion fluxes (Fig. 2F)

Our conceptual framework integrates three structural levels of the organism – the somatic cell structure, the stem cell structure, and a global (aggregate) cell structure (Fig. 2A)- and their mechanisms of interaction needed to restore anatomical and bioelectric homeostasis. The stem cell network is the engine of regeneration (Fig. 2A Level 2 & Fig. 2B). Somatic cells form networks with local neighbourhood communications (Fig. 2C), represented by perceptron neural networks, with local interactions in each of the three tissues (head, body, tail) of the organism. Stem cells form a similar, but one body-wide, perceptron network. In both networks, perceptrons communicate their bioelectric state with their neighbours using three simple generic communication motifs. These motifs were trained to represent the neighbourhood rules applicable to the three possible locations of a cell (interior, corner, or border) in the respective networks. They identify the presence or absence of neighbours and thus simplify body computing. The two perceptron networks identify missing neighbours based on changes in the neighbour’s bioelectric state. Global nodes form a 13-node network represented by an Associative Memory Neural Network (AMN) (a modified form of a Hopfield network) (Fig. 2A - Level-3 & D) that recognises and restores the body-wide homeostasis bioelectric pattern of the nodes after any perturbation or damage. These nodes consist of a cluster of stem cells surrounded by somatic cells (Fig. 2E) that together constitute the organism. The AMN is trained to store the homeostasis bioelectric pattern in its attractor. Further, an Information Field containing a minimal body plan for the tissues (tissue length (d), the aspect ratio (length/width), the number of corners (n)) and the homeostasis bioelectric state surrounds the organism and is accessed by stem cells and the AMN. Fig. 2G shows the two main axes of the organism, Anterior-Posterior (A/P) and Dorsal-Ventral (D/V), respectively.

The Methods Section provides the organisational and functional description of the framework that features five stages (Fig. 7A & B) of regeneration. These stages cover how somatic and stem cell networks detect a bioelectric state change anywhere in the system through the changes in their neighbour’s bioelectric state; how perceptron communication then identifies whether the change is due to normal function or damage; and, in the case of no damage, how the AMN restores bioelectric homeostasis; and, in the case of damage, how perceptron communication identifies the missing neighbours in the somatic and stem cell networks depending on the damage; how neighbour stem cells of damaged stem cells regenerate new stem cells that migrate to the damage site in the somatic cell network and completely repair the damage with the help of the somatic cell network; and how the stem cell network subsequently informs the AMN that incrementally restores bioelectric homeostasis by updating the nodal voltage until it matches the voltage pattern stored in its attractor. The Methods section also describes how the AMN, the stem cell, and the somatic cell networks are trained to accomplish these tasks.

## RESULTS

### Implementation of the framework for automated pattern regeneration and bioelectric restoration: Organism maintains body-wide immortality and bioelectric homeostasis

In this section, we demonstrate the implementation of the framework on a simple synthetic (*in silico*) worm to show that it efficiently and robustly recovers from both very simple to very complex damage. We consider two main types of damage: partial tissue damage with intact stem cells; and severe damage incurring the loss of stem cells and tissue segments or whole tissue (head or tail). We illustrate how the framework efficiently restores the bioelectric pattern due to normal perturbation or damage.

### The model worm and its original geometric pattern and bioelectric state

The artificial model organism is a simple worm with head, body, and tail tissues consisting of 150 stem cells (red dots) distributed throughout the organism. There are 3750 somatic cells surrounding the stem cells (tiny dots which fill the whole organism) (Fig. 2A). The organism consists of repeated patterns (square blocks) of stem cell-centred somatic cell neighbourhoods, sized 5 × 5 (Fig. 2C) (24 somatic cells per stem cell). The arrows in Figs. 2B & E and Fig. 2C respectively show the closest neighbours of the stem cells and the somatic cells. The colours in Fig. 2A show the body-wide bioelectric gradient, indicating decreasing cell membrane voltage, from the head to the tail, and from the middle of the body to the border regions [7, 8]. The homeostasis bioelectric pattern is negative throughout the planarian indicating that the membrane voltage is hyperpolarised.

### Organism restores body-wide bioelectric pattern due to normal perturbation (without damage)

The AMN refers to body-wide associative memory. It is a mechanism that helps its nodes communicate, through links with trained connection weights, to collectively maintain the voltage in their constituent cells following any perturbation. The AMN is trained with Hebbian learning using a large number of perturbed voltage patterns of nodes, representing changes due to normal physiological function and damage. The trained network remembers the original bioelectric pattern in its attractor and stores it in the Information Field (see the Methods Section). Changes to nodal voltage triggers the AMN to restore the original bioelectric pattern.

In the resting condition, the original body-wide bioelectric pattern takes the form of Fig. 3A. When a cell or a few cells experience a change in voltage, the corresponding AMN node *k* that the cells reside in, also undergoes a change in its voltage *V*_*k*_ (Fig. 3B). This change triggers the AMN network to restore the original bioelectric pattern of the affected nodes and cells (Fig. 3F). This case follows Stages 2->5->1 of the regeneration framework (see Fig. 7B). These stages are: detection of bioelectric change and confirmation of no damage (Stage 2), restoration of the bioelectric state (Stage 5) and a return to monitoring (Stage 1). When somatic cell(s) experience a voltage perturbation, Stage 2 (Fig. 7B) is activated to determine if this change is due to normal functions or damage. Voltage changes due to normal physiological functions are typically below 10% [34]. The affected cells in the somatic cell network send bioelectric signals through the GJs and ion fluxes to the stem cells. Both the stem and somatic cell networks detect this change. The affected somatic and stem cells check for missing neighbours by applying the three perceptron communication motifs for the corner, border, interior cells (see the Methods Section and Figure S1). These perceptron motifs are trained to take the neighbours’ voltage as input and predict their presence or absence. In this case, the affected cells find that there are no missing somatic or stem cells, and the stem cell network instructs the AMN to restore the bioelectric state (Stage 5). Beginning with the altered voltage pattern as input, the AMN restores the original voltage pattern stored in the attractor in a series of steps. This process of gradual restoration allows cells to incrementally produce the required changes in voltage. The AMN remains active until the system has returned to the original bioelectric state (Fig. 3F) (Stage 1). Even if a greater number of (or all) cells experience deviations from the equilibrium voltage state (see Fig. 3C), the AMN similarly restores the original bioelectric pattern in a series of steps (Figs. 3D - F). The framework thus restores bioelectric homeostasis due to any physiological disturbance.

**Fig. 3.**
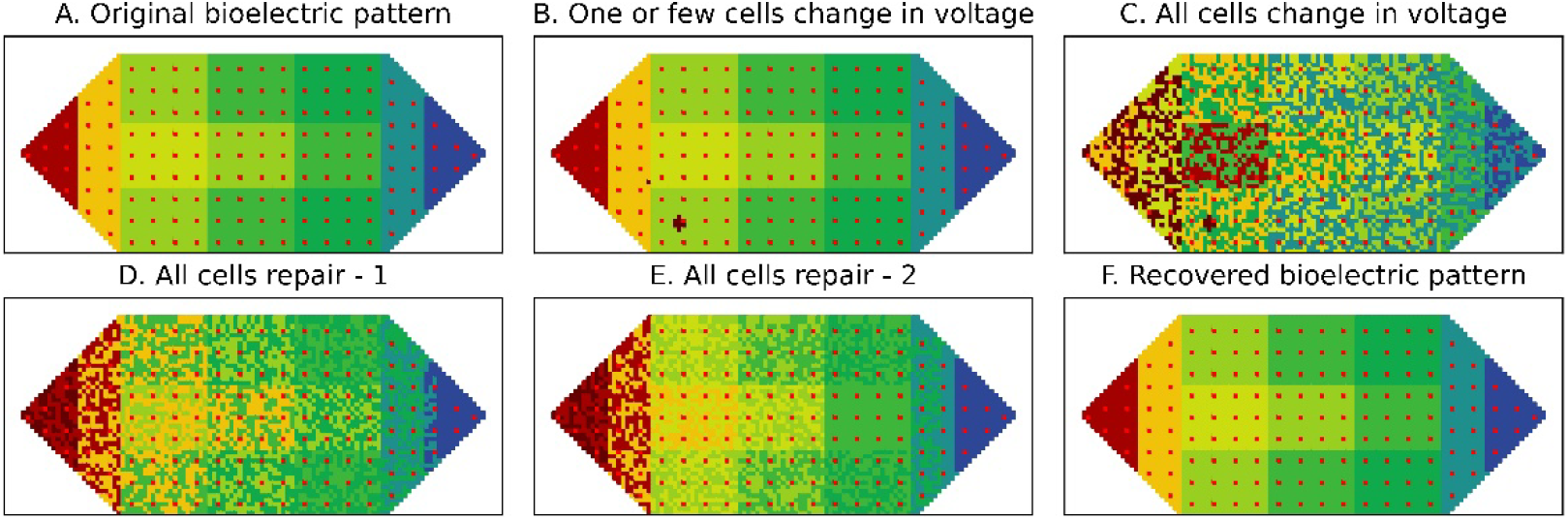
Perturbation and restoration of body-wide bioelectric voltage pattern under normal physiological function. (A) Normal equilibrium voltage pattern (homeostasis); (B) One or a few cells in a global node experience a voltage change - up to ±10%; (C) All or most of the cells in all nodes experience voltage change - up to ±10%; (D&E) Stages of recovery of the bioelectric state via activation of the AMN; (F) Recovered original voltage pattern.

### Organism recovers from small or large scale damage

#### Case 1: (a) Framework recovers from simple damage with one somatic cell missing

Here cell death occurs due to ageing or harmful factors such as trauma or toxic chemicals (Fig. 4A and enlarged in Fig. 4B). This case involves simple damage where only one somatic cell is missing anywhere in the organism. This damage case helps illustrate some fundamentals of the framework simply. When a somatic cell dies, connections between the damaged cell and its neighbour somatic cells through the GJs are lost. The neighbour cells’ voltage increases by >10% [34] due to the electrolytes released from the dead cell. This process activates Stages 2->3->4->5->1 of the framework (Fig. 7B). These stages are: detecting bioelectric change and confirming damage (Stage 2), identifying damage (Stage 3), restoring damage (Stage 4), restoring the bioelectric state (Stage 5) and returning to monitoring (Stage 1). The somatic cell network experiences the voltage change and sends bioelectric signals through the GJs and ion fluxes to the stem cells (Stage 2). After sensing the increased voltage, indicative of damage, the affected somatic cells apply the three perceptron communication motifs to confirm that damage has actually occurred (Stage 3). The cells with missing neighbours change their status to “damaged”. As illustrated in Fig. 4B, the “damaged” somatic cells (yellow cells) are neighbours of the damaged cell (the black cell) and become the damage border. The affected stem cell assesses the neighbouring stem cells using the relevant perceptron motifs and determines that all of its neighbour stem cells are intact. Here, only one somatic cell is lost.

**Fig. 4.**
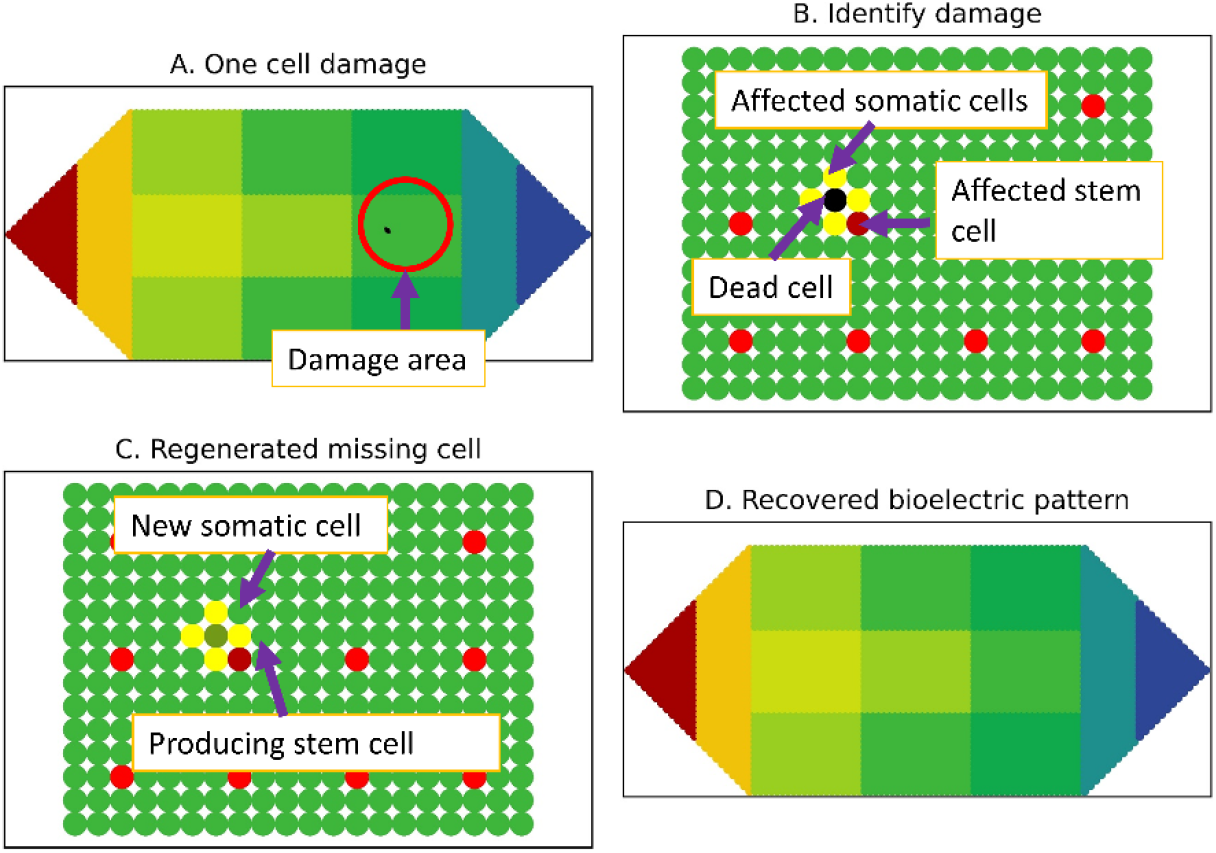
An example of a somatic cell damage and its regeneration and bioelectric state restoration. (A) One damaged somatic cell (black cell); Solid circle highlights the affected cell area; (B) Damaged region showing neighbours (yellow cells) of the damaged cell. The red cells are stem cells in this tissue; The dark red cell is the nearest stem cell affected by the damage; (C) Regeneration of the missing cell by the affected stem cell (the producing stem cell); the regenerated cell has the voltage of the producing stem cell; (D) AMN recovers the equilibrium bioelectric pattern.

In regeneration (Stage 4), the affected stem cell (the dark red cell in Fig. 4B) moves to the damage border and divides to produce a new somatic cell. The new cell’s voltage differs from that of the original cell and, in the absence of clear evidence, is assumed to be similar to the voltage of the stem cell that produced it (Fig. 4C); in reality it can either be this or between the original voltage and that of the somatic cells in the damage border. After regeneration, the new cell establishes connections with its neighbours and the relevant neighbourhood rules and corresponding communication motifs. All cells in the damaged border re-check their neighbours and update their status. As all these cells now have the correct number of neighbours as in the initial state, their status returns to “normal.” The stem cell returns to its original place in the tissue and activates the AMN to restore the bioelectric state (Stage 5), similar to the case without damage. The increased voltage in the cells in the damaged area (yellow and dark red cells in Fig. 4C) results in a slight increase in the corresponding AMN node’s voltage. Upon activation, the AMN uses the bioelectric state after regeneration as input and, in a series of steps, returns the altered nodes’ voltage to the normal state (Fig. 4D).

#### Case 1: (b) Framework recovers from damage, with multiple missing somatic cells

Successful regeneration and restoration of bioelectric homeostasis after damage to multiple somatic cells (Section S3, Case 1 in Supplementary Materials).

#### Case 2: (a) Organism recovers from damage involving a stem cell and its surrounding tissue

This example is a more complex damage case where a stem cell and a considerable number of somatic cells (about 24 cells) are gone (see Fig. 5A). The damaged region is highlighted in Fig. 5B (black cells) to show the missing stem cell and its surrounding tissue. As in Case 1, somatic cells neighbouring the damaged cells first detect the bioelectric change and alert the stem cell network (Stage 2). Since the voltage change is greater than 10%, they attempt to identify the damage (Stage 3). Neighbours of the damaged cells establish the somatic cell damage border (the yellow border in Fig. 5B) using the relevant perceptron motif (in this case, for an interior cell). Fig. 5B shows that the four nearest stem cells (the dark red cells) experience an altered bioelectric state and they check their stem cell neighbours using the relevant perceptron motif (for interior cells). These four stem cells establish the stem cell damage border and identify that one neighbour stem cell is missing. To identify the nature and size of damage, three pattern primitives (Methods Section) are applied to the stem cell damage border to determine if the damage is enclosed, has open borders, or whole tissues are missing.

**Fig. 5.**
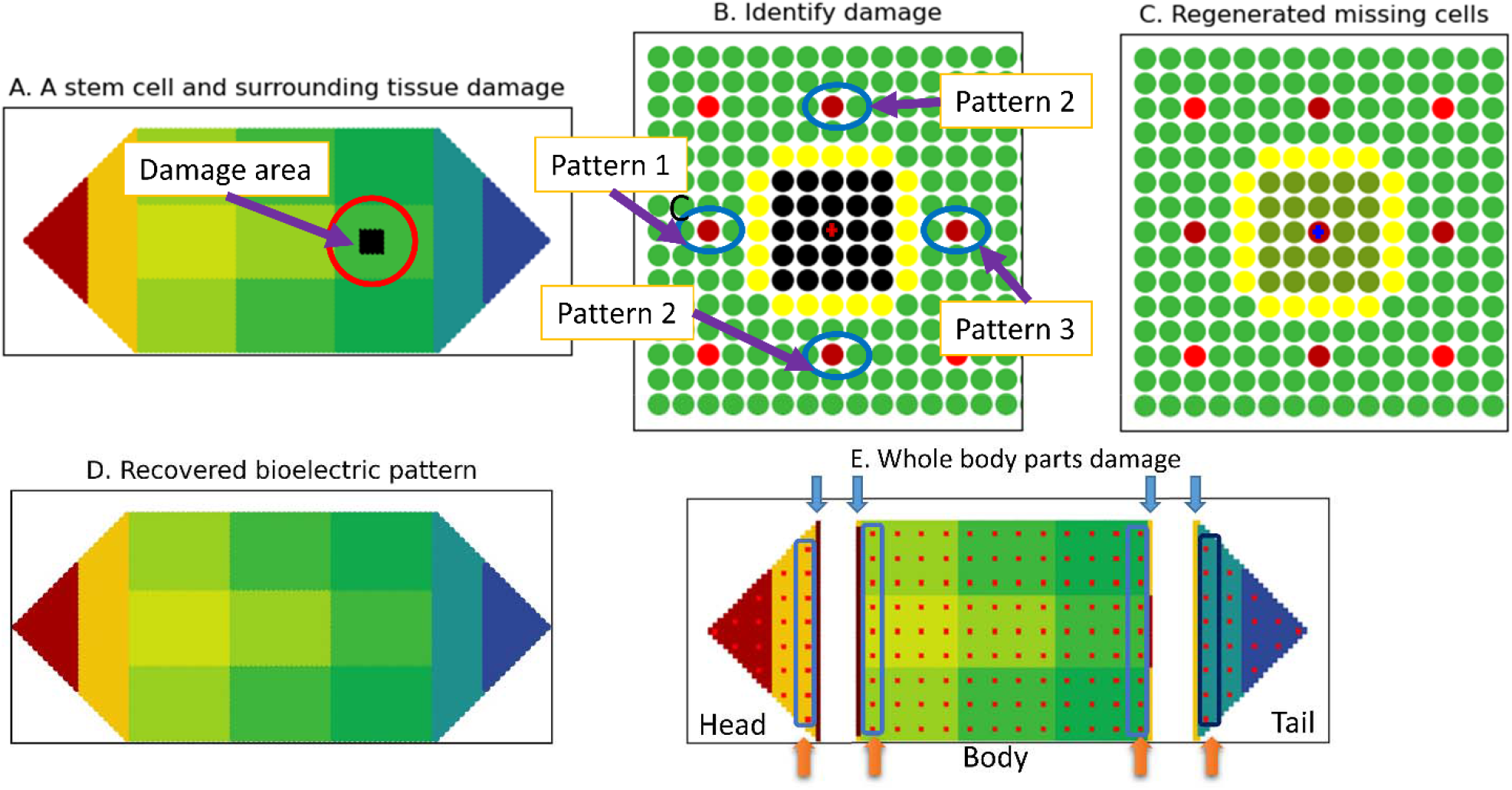
Damage involving a stem cell and surrounding tissue and its regeneration and bioelectric state restoration. (A) A stem cell and surrounding tissue damage (black cells); (B) Identification of damage - yellow cells show the damage border of the somatic cell network. The dark red dots indicate the damage border of the stem cell network. The middle black cell with a red cross in the damage region indicates the missing stem cell; (C)Regenerated missing stem cell (red cell with a blue cross) and somatic cells and the bioelectric state after regeneration; (D) The AMN restores the body-wide bioelectric pattern; (E) Damage to whole body parts – the worm is completely cut into three separate tissues: head, body, and tail. Blue arrows indicate the somatic cell damage borders identified by the respective tissue somatic cell networks in the three tissues. Orange arrows indicate the stem cell damage borders identified by the stem cell network.

The three primitives are basic patterns that help identify the nature of the stem cell damage by assessing if there are border stem cells (i) to the left, (ii) above or below, or (iii) to the right of the damage, representing pattern primitives 1, 2, and 3, respectively (see Fig. 5B). For example, if all patterns apply, the damage is local (encased or with an open border). If only some of the patterns apply, there is large scale damage; for example, if only one pattern applies, this indicates complete tissue damage. In Fig. 5B, all three patterns apply, as there are border stem cells to the left, right, and above and below the damage, indicating local encased damage. To regenerate (Stage 4), any stem cell in the border can migrate to the (yellow) somatic cell damage border and begin the regeneration process. The stem cell first produces a new stem cell with the same voltage as itself. The new stem cell then produces new tissue with 24 somatic cells, guided by the somatic cell damage border, to achieve accurate regeneration. In this case of interior damage, new somatic cells fill the (black) damage region, at which point regeneration stops (Fig. 5C). To restore voltage (Stage 5), the stem cell network activates the AMN to revert to the original pattern (Fig.5D).

#### Case 2: (b) Organism recovers from damage involving multiple stem cells and surrounding tissue

Successful regeneration and restoration of bioelectric homeostasis after damage, involving multiple stem cells and surrounding tissue (Section 3 Case 2 in Supplementary Materials).

#### Case 3: (a) Organism recovers from damage to whole body parts – completely missing head, body, or tail

##### Sensing damage and identification of lost tissue/s

In this last example, the worm is cut into three parts: head, body, and tail (Fig.5E). As described in the previous examples, stem cells and somatic cells in the severed parts sense the voltage change (Stage 2). First, somatic cells identify the border of the tissue damage (blue arrows in Fig. 5E) using perceptron motifs. The affected stem cells check their neighbour stem cells using the perceptron motifs. This process establishes the border of stem cells in each tissue; specifically, four stem cell borders (one each for the head and tail and two for the body) (orange arrows in Fig. 5E). The stem cells identify the severity of damage (Stage 3). Applying the three pattern primitives to the actual stem cell borders reveals that these border stem cells encompass only one generic pattern primitive (i.e., border stem cells found either to the left or right of damage), which means whole tissue damage. Border stem cells then identify the type of missing tissue (head, body, or tail) by applying two specific rules (Methods & Section S1.3 in Supplementary Materials) that recognise the direction of disrupted bioelectric communication in the worm:

###### Severed head tissue

The border stem cells in the head tissue do not receive any signals along the AP direction (from the body side). They still receive normal signals (from the head side), so they recognise that the body is lost.

###### Severed body tissue

For the damaged side on the left: stem cells do not receive signals along the AP direction (from the head side). They receive signals from the tail side; therefore, they recognise that the head is lost. Similarly, for the damaged side on the right: stem cells do not receive signals along the AP direction (from the tail side). They still receive normal signals (from the head side), so they recognise that the tail is lost. This indicates that the body tissue is separated.

###### Severed tail tissue

The stem cells nearest to the damage border identify the type of damage as the head and body missing: these stem cells do not receive signals along the AP direction (from the head side). They still receive normal signals (from the tail side). All three separated tissues regenerate (Stage 4) into three separate worms. The case of the head regenerating the whole worm is described below.

##### Regeneration: Head regenerates body and tail

Stem cells in the damaged border (Fig. 6A) produce new stem cells for the body and the tail, and transfer respective minimal tissue shape information from the Information Field (AR, d, n) to them. They then produce their respective tissues (body and tail) concurrently (Fig. 6B). The new cell’s voltage is lower than the producing stem cells due to the bioelectric gradient from head to tail. The small body and tail grow incrementally into the exact original form following the shape information obtained from the Information Field (Figs. 6B-E). The bioelectric pattern (soon after regeneration) is shown in Fig. 6E. Finally, the AMN restores its network configuration and retrains the weights using the original bioelectric pattern stored in the Information Field. This way, it restores the repaired nodes’ voltage (Stage 5) and brings the whole worm back to the equilibrium voltage pattern (Fig. 6F) (Stage 1). The severed body and tail also regenerate the whole worm using the same process.

**Fig. 6.**
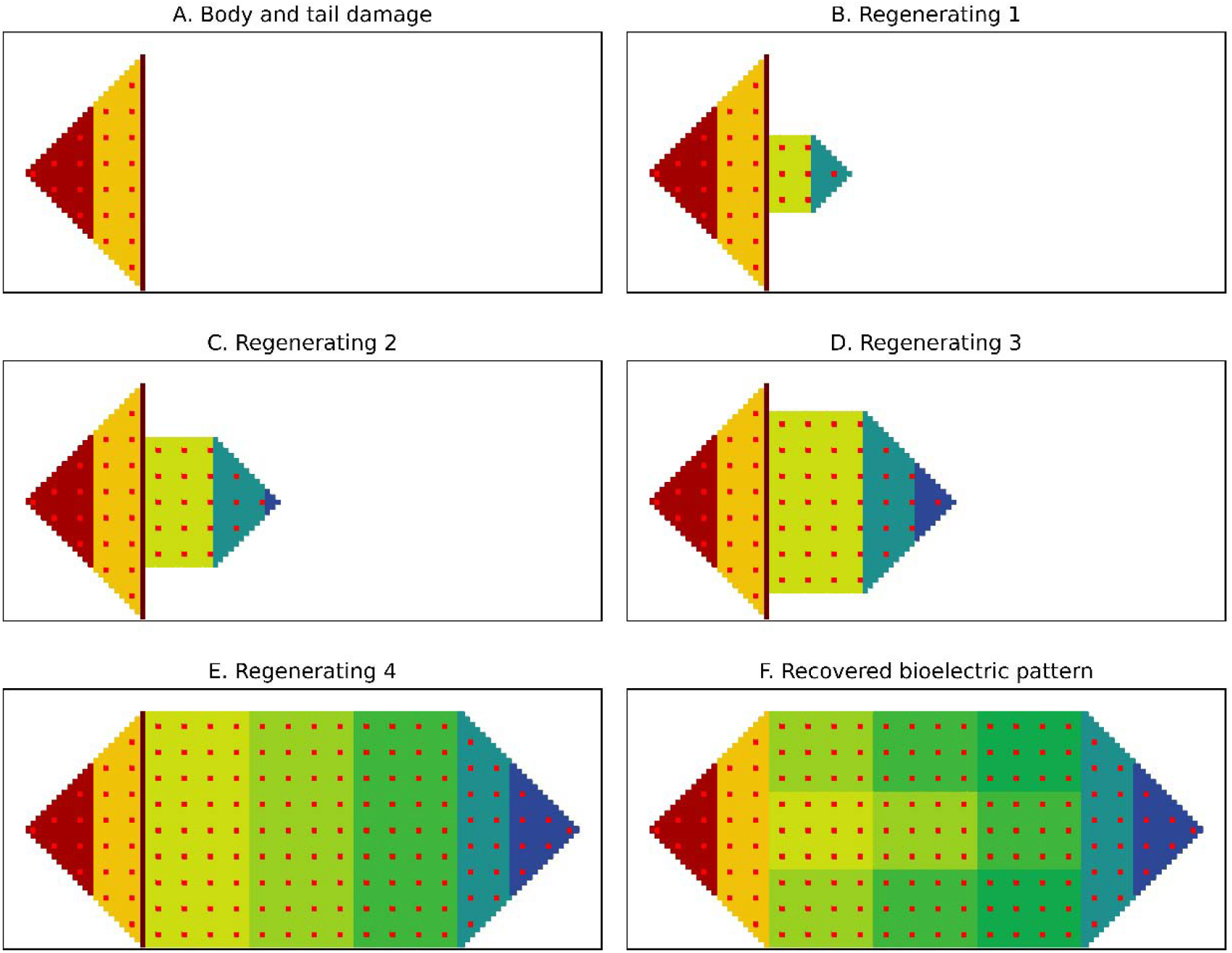
The head regenerates the body and tail to restore the whole form and bioelectric pattern. (A) Severed head tissue showing the stem cell border close to the damage site; (B-E) Regeneration of the body and tail stem cells by the stem cells in the stem cell border and concurrent regeneration of head and tail tissues until full recovery of anatomy and (F) AMN restores the body-wide bioelectric pattern.

#### Case 3: (b) Organism recovers from damage to whole tissue along with interior damage

Successful regeneration and the restoration of bioelectric homeostasis after this type of complex damage is presented in Section S3 Case 3 in Supplementary Materials.

#### Case 4: Organism recovers from more complex damage where worm is split into two with vertical and horizontal cuts

Successful regeneration and the restoration of bioelectric homeostasis after this type of complex damage is presented in Section S3 Case 4 in Supplementary Materials.

## DISCUSSION

In this study, we asked whether it is possible to conceptualise the mechanisms and algorithms of regeneration of an amazing model organism like a planarian. These organisms achieve body-wide immortality and maintain bioelectric homeostasis which enable them to continue functioning perpetually. To answer this question, we proposed an autonomous computational soft robotics self-repair framework that displays body-wide immortality and restores bioelectric homeostasis, much like the planarian, in a simple *in silico* worm. We demonstrated the application of the framework to successfully detect and completely and accurately repair diverse forms of damage and restore bioelectric homeostasis. Our framework is an intelligent control system within a neural computing paradigm for collective cellular decision-making for the dynamic restoration of morphology and physiology. The model combines three levels: somatic cell networks and a stem cell network, both represented as locally interacting perceptron networks for damage identification and full and accurate recovery from diverse forms of injuries, and a high-level AMN, for the restoration of body-wide bioelectric homeostasis. We demonstrated the efficacy in recovery using several very small to very complex damage cases through implementation of the framework. The three networks function collectively to maintain the anatomical pattern and bioelectric homeostasis. We believe that this is the first work that provides a conceptual basis and algorithms explaining how an organism like the planarian can maintain and restore form and function.

In this framework, bioelectricity is a fundamental property of the cells. It helps cells communicate with each other to maintain the original form and body-wide bioelectric gradient. The somatic cell networks transfer bioelectric signals to the stem cells and identify the border of tissue damage that guides the stem cells to injury location. As a body-wide network, stem cells are the engine of regeneration. As a result of bioelectric signalling between somatic and stem cells, stem cells can detect small local damage or complex large-scale damage involving lost stem cells or whole tissues and repair the damage completely and accurately with the help of the somatic cell networks. The AMN mimics an unknown mechanism in planaria that maintains the body-wide bioelectric gradient, along and across the worm.

Our study contributes a comprehensive soft robotics framework to generate hypotheses about the mechanisms of regeneration to advance the knowledge in this field. It also contributes several concepts and algorithms to significantly advance computational models of regeneration involving minimal computation. We compare our model to the state-of-the-art regeneration models to demonstrate its novelty and efficacy.

Our previous regeneration framework [26] is the closest to the current work. The previous framework regenerated an organism of three tissues using binary communication between cells without using bioelectricity. We have greatly improved our previous study. Firstly, we have proposed a greater number of stem cells, about 4% of the total, distributed within tissues. In the previous framework, only one stem cell represented each of the three tissues. Secondly, we introduced two types of bioelectric communications-through Gap Junctions (GJ) and ionic fluxes. Somatic cells use physical GJs to connect to, and communicate with, adjacent cells. Stem cells maintain long-range bioelectric communications propagated through the intermediate somatic cells (ion fluxes moving through GJs and then released into the surrounding environment). While stem cell communications are contained within a local region, minimising computation, the stem cell network is distributed throughout the organism. The current model is more biologically realistic compared to our previous one [26] where we used direct, long-range interactions by which the stem cell in the head can communicate with the stem cell in the tail. Thirdly, we used an AMN as a mechanism to help the worm maintain the bioelectric gradient in an equilibrium state. We found an intriguing minimal network configuration for the AMN (with tail to head inhibition and head to tail activation) that efficiently stores and retrieves the equilibrium bioelectric pattern of 13 macro (global) nodes, each consisting of approximately 300 somatic cells and 12 stem cells (Section S1.4, Supplementary Materials). This discovered network configuration points to the possible existence of simplified and directional networks as opposed to fully connected networks within cellular structures that perform complex tasks such as the restoration of body-wide homeostasis. This structure helps the worm maintain the bioelectric gradient in A/P and D/V directions. We established similar directional activation and inhibition in our configuration of the stem cell network to conform to the AMN (Fig. 2B).

Finally, we used three perceptron motifs which are trained for the stem and somatic cell networks so that they can determine the missing neighbour stem and somatic cells. For the stem cell network, which is an organism-wide perceptron network, the weights of these motifs represent the effective conductance between two stem cells, and correspond to the conductance of the GJ of intermediate somatic cells and ion fluxes in living multicellular structures. For the tissue specific somatic cell networks, the weights of these three motifs represent the bioelectrical conductance of the GJs between somatic cells. In the previous framework, we represented stem cells using a linear neuron with fixed weights, and a somatic cell using a perceptron with fixed weights (equal to 1). In the new framework, the three motifs use cell voltage as inputs to communicate output for a somatic or stem cell located anywhere in the organism through trained network weights. The two cell networks collaborate to repair damage. The AMN then restores the bioelectric gradient. To identify various types of stem cell damage, ranging from one or a few stem cells to completely missing whole tissues, we proposed three simple pattern primitives applied to the stem cell damage border. In the case of missing whole tissues, we additionally proposed two simple rules involving the direction of the disrupted communication flow, from the head to tail or vice versa, to identify the exact missing tissue.

Other prior models have attempted single or multiple tissue regeneration and have achieved varying degrees of success via a large amount of computation. None has achieved complete and accurate regeneration or presented a comprehensive framework. None has attempted bioelectric restoration. Bessonov et al.’s [13] tissue regeneration model assumes that an individual cell communicates with all other cells and remember the total signal intensity value it receives and some regeneration rules. Regeneration is based on restoring the total signal intensity of each cell. However, there is a high computational burden due to the extensive communication between cells. The model also has limited success in accurate recovery. Due to the simple local communications, the computational burden in our model has been drastically reduced, while achieving complete and accurate recovery. In Tosenberger et al.’s [14] circular tissue model, regeneration is accomplished via stem cells that are not damaged. Our framework applies to multiple tissue shapes and allows stem cell damage. The stem cell network replaces missing stem cells. In their model, tissue regeneration requires all cells to compute the received signal intensity and compare it to a threshold value, which is computationally burdensome. Further, it assumes that the tissue has a survival region and that cells produced outside of this region are killed to achieve the required shape, which is biologically unrealistic. In our framework, local communication in somatic perceptron networks in tissues identifies the damage boundary which guides complete and accurate recovery. In extreme cases involving tissue loss, minimum body plan information is accessed from the Information Field.

De et al.’s [4] model is one of the more biorealistic models. It involves interactions between a neural network (representing the organism’s nervous system), and non-neural cells (representing tissue), to recover from damage using physiological variables. It has achieved reasonable regeneration success (55%–80%), depending on damage severity. However, it does not recognise damage and generates and kills many cells before achieving a partially recovered form. Our framework detects damage and accurately recovers the complete form using simpler and more efficient computations in stem and somatic cell networks. As long as a single stem cell exists, our model can regenerate the whole organism correctly. Ferreira et al.’s [17] agent-based models require much communication between cells to discover cell structure and store much data. Further, these models cannot regenerate larger sized tissues as information can only go a limited distance or signals decay with distance. By contrast, the information/data our framework carries in regeneration is high-level and generic, with three simple perceptron communication motifs. It contains few rules relating to the stem cell damage border pattern and a minimum amount of information on shape to achieve efficient regeneration in any sized tissue. Importantly, it involves minimum computation.

Although our autonomous framework has emulated important aspects of planarian regeneration, it has a number of limitations that can be improved. First, the form of the chosen organism is simple and needs to be extended to a more realistic shape like the real planarian or other organisms. Second, in each AMN node, the somatic cells’ voltage is assumed to be constant. These reference voltage values could be made to vary within a node to achieve a more refined bioelectric gradient. The chosen voltage pattern represents that which can be simulated by partial differential equation-based models of voltage patterns produced by bioelectric communications between adjacent cells through GJs (as in [35]). Our framework can integrate these bioelectric models and the models of diffusion of important signalling molecules with molecular signalling pathways to activate biological processes (such as cell division in regeneration), to represent both molecular and cellular levels of regeneration. These additions will enhance the framework’s collective intelligence and decision-making capacity by capturing fundamental processes involved in regeneration, meaning that the system more accurately resembles living planaria and living systems.

## METHODS

We present the framework in two parts: the *conceptual basis* and *functional description*. In part one, we elaborate on the framework’s conceptual basis and its organisation (Fig. 7A). Part 2 provides a functional description of the framework (Fig. 7B), with algorithmic/computational structures for a complete and accurate restoration of body patterns and bioelectric homeostasis.

**Fig. 7.**
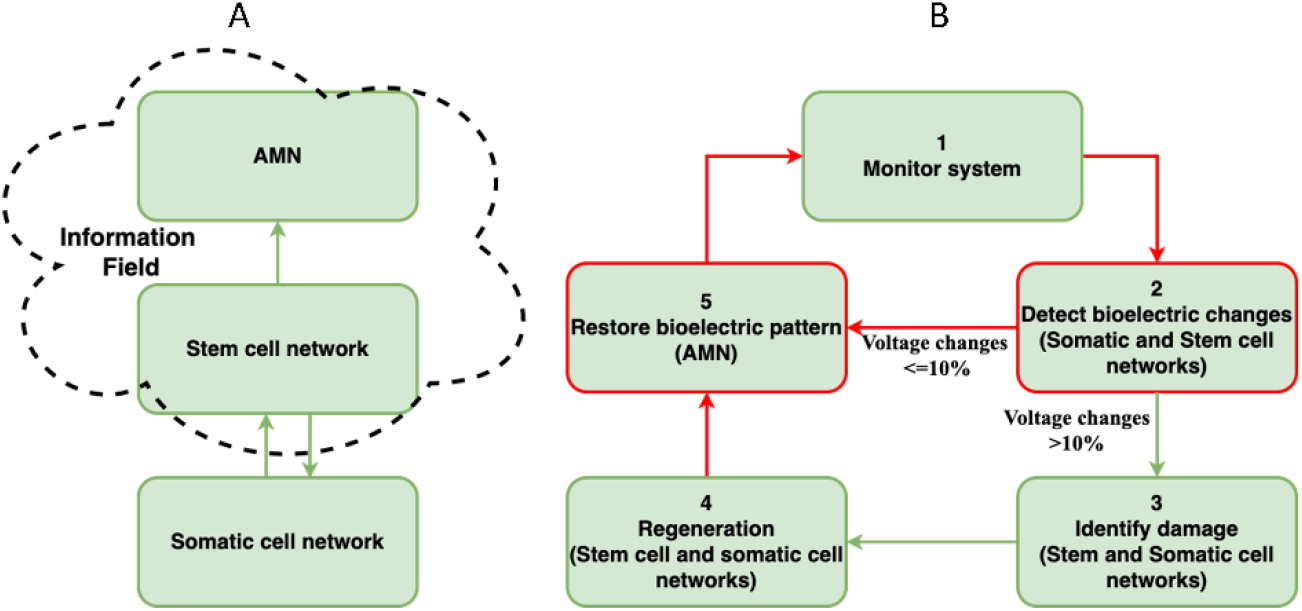
Overview of the framework for anatomical and bioelectric homeostasis. (A) Conceptual basis involving organisation of three networks and an Information Field. (B) Functional aspects of the framework involving five stages for restoration of both form and function.

### 1. Conceptual basis and organisational view of the self-repair soft robotics framework

We introduce three types of cellular networks in our framework that communicate using bioelectricity to restore both pattern and bioelectric homeostasis (Fig. 7A): the somatic cell networks, the stem cell network and the AMN that operate as an integrated whole. Specifically, the framework proposes a high-level (macro-level) body-wide AMN as a mechanism for maintaining and restoring voltage homeostasis (Level 3 in Fig. 2). This is a recurrent neural network, trained to retain the body-wide bioelectric gradient in its attractor state and recover it from normal perturbations and after damage recovery. The AMN’s purpose is to: (1) produce a simple model that maintains the body-wide bioelectric pattern; and (2) be a flexible and fault-tolerant system without excessive micromanagement of the bioelectric state (membrane voltage of cells). As there are approximately 3750 cells, we grouped the cells into clusters, each consisting of a group of stem and somatic tissue cells. We used clusters as AMN nodes (Fig. 2E).

For pattern regeneration, the framework employs somatic cell networks (one for each tissue) (Fig. 2C) and a stem cell network distributed throughout the organism (Fig. 2B). These networks are locally interacting perceptron networks. The stem cell network is the driver or engine of regeneration. We design regeneration mechanisms through bioelectric communication between a large number of stem cells and somatic tissue cells (Fig. 2F). Although voltage varies along and across the organism, there are only three patterns of perceptron communication (interior, border and corner cells) within the somatic cell networks and the stem cell network. The local interactions and three communication motifs greatly minimise the amount of computation. We also include an Information Field where the stem cells store a minimal body plan (a few basic tissue dimensions) for recovery from extreme damage, where all exterior boundaries are gone or where there is no trace of the original pattern boundary in the damaged tissue. In our simple worm, the Field stores the following pattern information for the three tissues: the length of the body tissue (d), the aspect ratio (AR) (the length to width ratio) of the head, body, and tail (AR=1, 3, 1), and the number (n) of corners of the head, body, and tail (n=3, 4, 3). All stem cells can access the Information Field. It also contains the AMN attractor that corresponds to the homeostasis bioelectric pattern. We present the algorithms, and the training and operation of these networks in part 2 of the description.

We retain some features of our previous framework [26] which only regenerated the form (not the function), of the organism. While we retain the perceptron configuration of the somatic cell network, the cell states are derived from actual voltages. Communication is now conducted through bioelectricity instead of binary codes [0,1], and perceptron motifs have trained weights instead of fixed weights. The stem cell network, also represented by a perceptron network, is totally new and much larger, with 375 stem cells distributed throughout the body, instead of the three stem cells represented by fixed linear neurons in the previous framework. Stem cell perceptrons have the same three communication motifs as their somatic cell counterparts. The AMN and the restoration of bioelectric homeostasis is novel. The AMN was trained using a large number of perturbed voltage patterns from the original homeostasis pattern to store the original pattern in its attractor. While the Information Field is similar, it has been expanded to contain the AMN attractor (the homeostatic bioelectric state). The Field can now be accessed by all stem cells and the AMN. Due to the presence of a larger number of stem cells and bioelectric communication, the operating system differs from the previous framework: it seamlessly integrates patterns and bioelectric homeostasis.

### 2. Functional overview of the framework – Collective intelligent decision making for autonomous form and function regeneration

The three neural networks operate collectively. They detect the size and location of the damage and repair it and subsequently restore the bioelectric state as presented in the high-level functional view of the system (Fig. 7B). The framework’s logic is highlighted in five stages that operate in two modes. The first mode involves restoring bioelectric homeostasis under normal physiological fluctuations in an undamaged organism where the system monitors (Stage 1), detects bioelectric changes in the cells (Stage 2), and if the change is smaller than 10% [34] indicative of no damage, it restores the bioelectric state using the AMN (Stage 5). The second mode involves damage repair followed by the restoration of bioelectric homeostasis. If the voltage change is greater than the 10% threshold indicative of damage, the framework activates the stem cell network and the affected somatic cell network(s) to determine the damage to respective networks (Stage 3). In Stage 4, the stem cell and somatic cell networks collaborate to repair the damage. In Stage 5, the AMN restores the bioelectric pattern. All three of the system’s networks sense changes in the bioelectric state that trigger one of these two modes of operation. Below, we elaborate on these modes of operation before describing the algorithms of the regeneration frameworks.

#### a) Bioelectric restoration under normal physiological function (stages 2, 5, 1)

When there is a perturbation in the bioelectric state, the system’s first task is to check if the voltage change is due to variations under normal physiological function or damage (Stage 2). In Stage 2, the somatic cells that have experienced a change in their voltage send this information to other cells through the GJs and the surrounding environment (ion fluxes). The stem cells receive this information and decide whether the change is due to damage or not using the ±10% voltage threshold. When it is below 10%, the stem cell network informs the AMN. The AMN works autonomously to restore the organisms’ homeostasis bioelectric pattern by iteratively reaching the stored attractor pattern (Stage 5).

#### b) Anatomical pattern and bioelectric restoration under damage (Stages 2, 3, 4, 5, 1)

When there is damage (indicated by a ≥10% voltage change), the stem cell network informs the AMN of this (Stage 2). The AMN waits while the affected stem cells check the damage (Stage 3) by applying the three perceptron motifs and identifying the stem cell damage border. Affected somatic cells check for missing neighbours and identify the tissue damage border. The somatic cells and the stem cells cooperate to recover the whole pattern from any damage (Stage 4). After regenerating, the stem cell network reactivates the AMN to restore the repaired cells’ and the organism’s voltage (Stage 5). The organism returns to normal anatomical and bioelectric homeostasis (Stage 1).

## Algorithms of regeneration

The framework consists of three levels, each with its own computational algorithm: the somatic cell network (Level 1), the stem cell network (Level 2) and the associative memory network (Level 3). Table S1 in the Supplementary Materials presents the three cell networks, their structure, properties, and activities.

### Level 1: Somatic cell network (perceptron network with local communication)

The somatic cell network recognises the intact pattern and damaged state of individual tissues through local bioelectric communication applied to three *communication motifs* (Figure S1 and Table S1, Column 2). These motifs specify the rules for the identification of neighbours of a somatic cell (inputs and the corresponding output state of perceptrons). Each somatic cell is represented by a perceptron that has two, three, or four neighbours depending on its location in the somatic cell network (interior, border, or corner, respectively). Therefore, a somatic cell receives two, three, or four inputs from its neighbours. This results in three communication motifs in the somatic cell network (Figure S1 and Eqns. S1 to S3 in Supplementary Materials). Table S1 provides the generic format of these motifs. We call these motifs because they can be applied to all somatic cells (corner, border, and interior), regardless of where they are located in the organism. As voltage varies throughout the body, only the inputs need to be standardised to 0 or 1 before presenting to the corresponding motif. As the voltage is negative throughout the system (Fig. 2A), 1 indicates the presence of a negative voltage and 0 the absence of any voltage. The output of the perceptron motifs is 0 or 1, indicative of the absence of one or more neighbours or the presence of all neighbours.

Somatic cells learn to recognise their neighbours dynamically, through trained perceptrons. For training, each perceptron motif starts with random values for the bias and input weights (Figure S1). These weights reflect the normalised electrical conductance of the GJs between somatic cells in a real planarian tissue. Actual conductance of the GJs for the whole organism are not currently known. These model conductance values are normalised representations obtained through training. Bias weight stabilises the outputs and represents any influence not captured by the inputs. Inputs 0/1 represent the presence or absence of somatic cells based on their voltage. The motifs were trained using Hebbian learning until convergence to a stable state, such that each motif can determine whether a neighbour(s) is present or not. The training was quick and successfully completed (see Table S2 for the three motif weights).

#### Identification of tissue damage by somatic cell networks

In the damage state, individual somatic cells, which are neighbours of damaged cells, sense the damage through a change in bioelectricity. They identify damage through the corresponding perceptron motif. The set of somatic cells, which recognise the missing neighbours, become the damage border. Thus, the somatic cell network can identify borders of any damage (Fig. 5A & B shows tissue damage using a black square that includes a missing stem cell). The tissue cells next to the damage receive an increase in voltage (yellow cells surrounding the back square (Fig. 5B)). Applying the motifs (in this case for interior cells), these tissue cells in the somatic cell network identify that their neighbours are missing and become the damage border. The nearest stem cells sense the border somatic cells’ elevated voltage and check if this is due to normal perturbation or damage.

### Level 2: Stem cell network – Operation of the regeneration engine (perceptron network with local communication)

The stem cell network (Fig. 2B) is represented by a locally interacting perceptron network. While it is similar to the somatic cell network, it is organism-wide and activated by sensed voltage signals, communicated through somatic cells. Further, these stem cell communications are either activation (+) or inhibition (-) signals depending on their position. From posterior to anterior (right to left), these cells inhibit neighbours. From other directions, they activate neighbours (Fig. 2B). This directed activation and inhibition pattern was extracted from the optimum pattern of nodal interactions found for the AMN network. This pattern is described in the next section, along with justification for the discovered pattern. As stem cells are within AMN nodes, stem cells were made to follow the AMN’s interaction format.

The stem cell network recognises its structure (and the damaged state) using the three perceptron motifs, with the same structure and weights as in the somatic cell network. The neighbours of the missing stem cells form the stem cell border. Depending on the received information, they restore either just the bioelectric state (no damage) or both damage and the bioelectric state (after damage).

#### Identification of stem cell and large-scale tissue damage

The stem cell network is the primary entity in damage detection and restoration. When stem cells sense increased voltage beyond the threshold, they first have to identify if there is stem cell damage, and if so, whether this is large scale global damage involving completely missing head or body tissues (Fig. 5E) or less destructive damage (local damage) involving one or more stem cells and associated tissue damage (Fig. 5A). Basically, the stem cells must identify the extent of damage to its network and determine the type of damage to the organism. To establish if stem cells have been damaged, this network applies the perceptron motifs to the cells that have experienced changes to their voltage status sensed through intermediate tissue cells. If all the affected stem cells report no missing neighbour stem cells, then the damage involves local tissue only, without stem cell damage. In this case, the somatic cell network identifies the border of the damage in the affected tissue(s) using the communication motifs. The nearest stem cell migrates to the damage border and repairs it. This damage does not involve tapping into the information field as relevant information is contained within the remaining tissue.

Although the forms of damage may be many, the stem cell network applies three pattern primitives to detect all types of stem cell damage. These patterns are based on the nature of the stem cell damage border created by neighbours of missing stem cells in the stem cell network. We explain these under two broad categories: local stem cell damage involving one or more stems cells (without whole tissue loss) (Fig. 5A); and large-scale damage involving the loss of whole tissues causing damage to large segments of stem cells (Fig. 5E). These categories are described in Section 1.3 of the Supplementary Materials.

In brief, the basic concept behind pattern primitives is very simple: all possible stem cell damage border patterns can be constructed using three simple pattern primitives (Fig. 5B). These three patterns refer to whether there are stem cells to the left, right and above or below the damage. In local (Fig. 5A) and open border damage, all three pattern primitives apply, as stem cells are found to the left, right, and above/below the damage (shown in blue circles in Fig. 5A). From this, it is recognised that the damage is local. In the case of large-scale tissue damage, only one pattern applies to the stem cell border around the damage. From this, it is recognised that the whole tissue(s) is missing. What exact tissue (the head, body etc.) is missing is identified by the direction of the flow of missing communication, whether it occurs from the head to the tail or vice versa (described in Section S1.3). For example, in the case of the severed head in Fig. 5E, the stem cell damage border in the head (indicated by the orange arrow) makes a simple pattern of one line of stem cells indicating the presence of stem cells to the left of the damage only. Therefore, only one pattern primitive applies which indicates whole tissue loss. Further, these damage border stem cells do not receive communication from the tail side which indicates that the body is missing. Similar logic applies to the severed tail that does not receive communication from the head side (refer to the respective orange arrow for the stem cell damage border in Fig. 5E). In the case of severed body tissue, there are two locations of damage, on the left and right sides. Each location of damage encompasses only one pattern primitive; that is, for the damage on the right hand side of the body, only pattern 1 applies, as the stem cells are to the left of the damage. For the damage on the right hand side of the body, only pattern 2 applies, as stem cells are to right of the damage, indicating whole tissue loss. Missing communication from both the head and tail sides indicates that the missing tissue is the body.

### Level-3: Global level Associative Memory Network (AMN) maintains and restores body-wide bioelectric homeostasis

This section proposes a hypothetical concept for dynamically maintaining bioelectric homeostasis in general functioning and restoring it after damage in an organism like the planarian. We ask, what minimal and generic approach could allow the remembrance and restoration of the bioelectric pattern represented by the membrane voltage of thousands of cells in the organism. We propose a global AMN, located on top of the stem cell network in our framework (Fig. 2A Level 3 & D). We assume that the body-wide bioelectric pattern is fault tolerant in that small-scale anomalies such as a failure in a single cell does not break the system. Equally, minor voltage fluctuations do not significantly alter the homeostasis voltage pattern. Considering the large number of cells in the organism, a more aggregate form of monitoring, at a macro level, that does not involve all the cells in the required computations is justifiable.

As proof of the concept, we divide the organism into 13 segments, each representing a group of stem and somatic cells with a corresponding level of membrane voltage (Fig. 8A). These constitutes the basic units or nodes of the AMN (Fig. 8B) involved in monitoring and restoring the body-wide bioelectric pattern. In this study, the voltage is assumed to be constant within an AMN node to keep it simple and test the AMN concept. This global level AMN is a recurrent neural network, which is a modified form of the fully connected Hopfield Neural Network (HNN) [36] that stores specific state vectors in one or a few attractors. We hope to find a network with minimum connectivity (as opposed to the full connectivity of all nodes in HNN) that is sufficient to remember the bioelectric pattern. The benefit of these networks in general is that for any distorted input vector (state vector), the network retrieves the original undistorted form of the same input vector. The AMN operates over the whole anatomical pattern of the organism and learns and memorises the body-wide bioelectric gradient (Fig. 8A). We provide a summary of the AMN development here (full details provided in Section 1.4 of the Supplementary Materials).

**Fig. 8.**
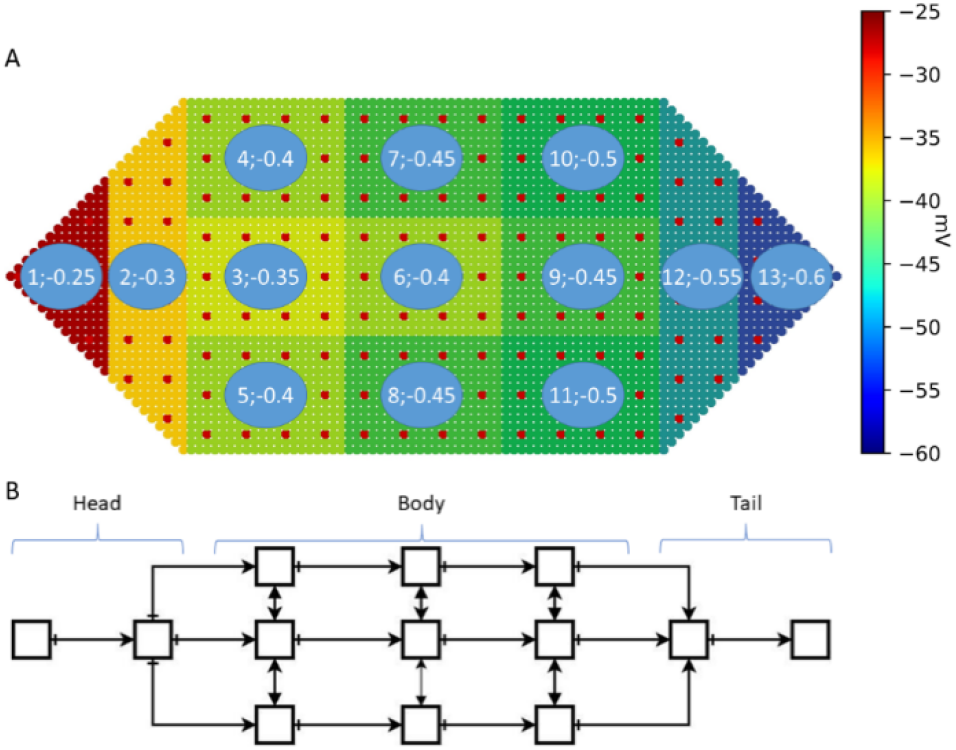
An organism with the original homeostasis bioelectric pattern and the AMN representing it. A) Innate body-wide bioelectric pattern and 13 tissue segments of the organism used to monitor it. The two numbers in each node are the node number and the corresponding membrane voltage (normalised). Colours indicate the magnitude of voltage. The whole voltage pattern across the organism denotes the homeostasis bioelectric pattern; B) The AMN, consisting of 13 segmented nodes, represents the whole organism and is designed to learn and remember the body-wide bioelectric pattern

The voltage pattern shown in Fig. 8A (i.e., 13 voltage values) is the pattern we wish to store in the AMN attractor. When this pattern changes due to normal perturbations or damage, the AMN should recall the original pattern and re-establish bioelectric homeostasis in collaboration with the stem and somatic cell networks. While we do not know if the planarian uses a mechanism similar to the AMN to keep a stable body-wide bioelectric pattern, we do know that the worm maintains a stable pattern. We assume that the approach it uses is flexible and fault tolerant, and that it achieves this aim without excessive micromanagement or extensive memory requirements.

Our proposed AMN has the following attributes. Each AMN node represents a cluster of cells: there are 300 tissue cells and 12 stem cells in a node (the head and tail nodes have a slightly different form to these cells). Each node has a constant voltage. The voltage of all the clusters form the body-wide bioelectric pattern. This arrangement allows the AMN nodes to sense voltage changes in their constituent stem and tissue cells in a global or average sense and react accordingly, to restore the bioelectric pattern under normal function or wait for the stem cell network to initiate repair after injury and complete regeneration. Within a node, neighbour somatic cells are connected by GJs with a particular conductance. Stem cells communicate through somatic cell GJs and ion fluxes. An AMN node containing these cells communicate with neighbour AMN nodes via pseudo GJ connections, reflecting effective or average conductance of the GJs connecting the somatic cells in the neighbour nodes (Fig. 2D). This system means that the AMN correctly resembles the cellular structure and interactions without losing information due to aggregation. It also helps it to accurately preserve the average membrane voltage profile throughout the body. These pseudo-GJ conductances are represented by the AMN weights that are determined by training. Training is performed through Hebbian learning, with a dataset generated by perturbing the original voltage pattern to represent both normal fluctuations and damage conditions.

Full details of the AMN, its training, and the final network configuration are presented in Section 1.4a of the Supplementary Materials. In brief, the model dynamics use asynchronous neuron state updates, with a rectilinear function (ReLU) in the AMN nodes (Fig. S2) that calculates the output (voltage) *s*_i_ of *i*^th^ node as follows (Eq. S5):

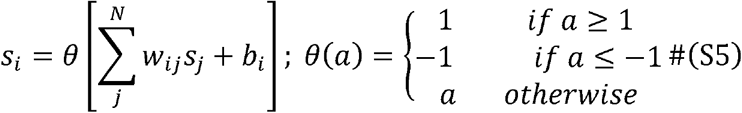

where *w*_*ij*_ is the connection weight between node *i* and *j, θ* is the transfer function (ReLU), and *b*_*i*_ is the bias weight of node *i*. To store the desired pattern, we set random values for the connection weights (w, b) and adjusted them using the Hebbian learning rule:

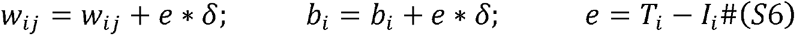

where *δ* is the learning rate, *e* is the difference between the target pattern *T* and the current input pattern *I*.

The proposed macro-level nodal configuration ensures the monitoring system is fault tolerant, and enables the AMN to represent the whole worm in a smaller representative network with grossly similar bioelectric properties to that of thousands of somatic cells. This configuration greatly simplifies the required computation needed to achieve the end result of maintaining homeostasis, where all somatic cells still contribute to the process. Moreover, by exploring sparse but meaningful connections between nodes in the AMN, we found a much-simplified network configuration that substantially reduces the computational burden on the network. This finding indicates the potential existence of efficient and high-level organism-wide networks for bioelectric control. Studies have shown that bioelectric perturbations can cause large scale morphological changes, such as the formation of a head in the tail position and a number of other severe anatomical distortions [10, 12].

Basically, with AMN, we do not eliminate anything but instead simplify the system. Since individual somatic cells are not nodes in the network, the network is flexible and fault tolerant. For example, the AMN will only be damaged if all cells in a node are damaged. Until then, the AMN collaborates with the stem cell network within the affected nodes to quickly restore bioelectric homeostasis after repairing any damage to cells within that node. When the perturbation is not due to damage, the network automatically restores the original bioelectric pattern. When a network node is damaged, the network’s spatial topology is broken. The network will lose the voltage value of the lost node and the corresponding connection weights. After regeneration, the connections from this node to others need to be re-established to restore the lost connection weights. After the nodes are repaired, the AMN retrains itself, by accessing the bioelectric pattern from the Field to create perturbed input training patterns, until the network learns to store the original pattern in the attractor and the connections and weights to the repaired nodes are re-established.

## Acknowledgments

The authors gratefully acknowledge the support of the following: S.S.- LURF Research Fund (New Zealand) and TNM - Doctoral Scholarship from VIED, Vietnam. They thank Prof Michael Levin at Tufts University for inspiration to do this work and Prof Don Kulasiri for reviewing and providing valuable comments on the manuscript.

## Supplementary Materials

### S1. Further details of the conceptual framework and methods

#### S1.1 Overall view of the components of the new framework and their attributes and properties

**Table S1.**
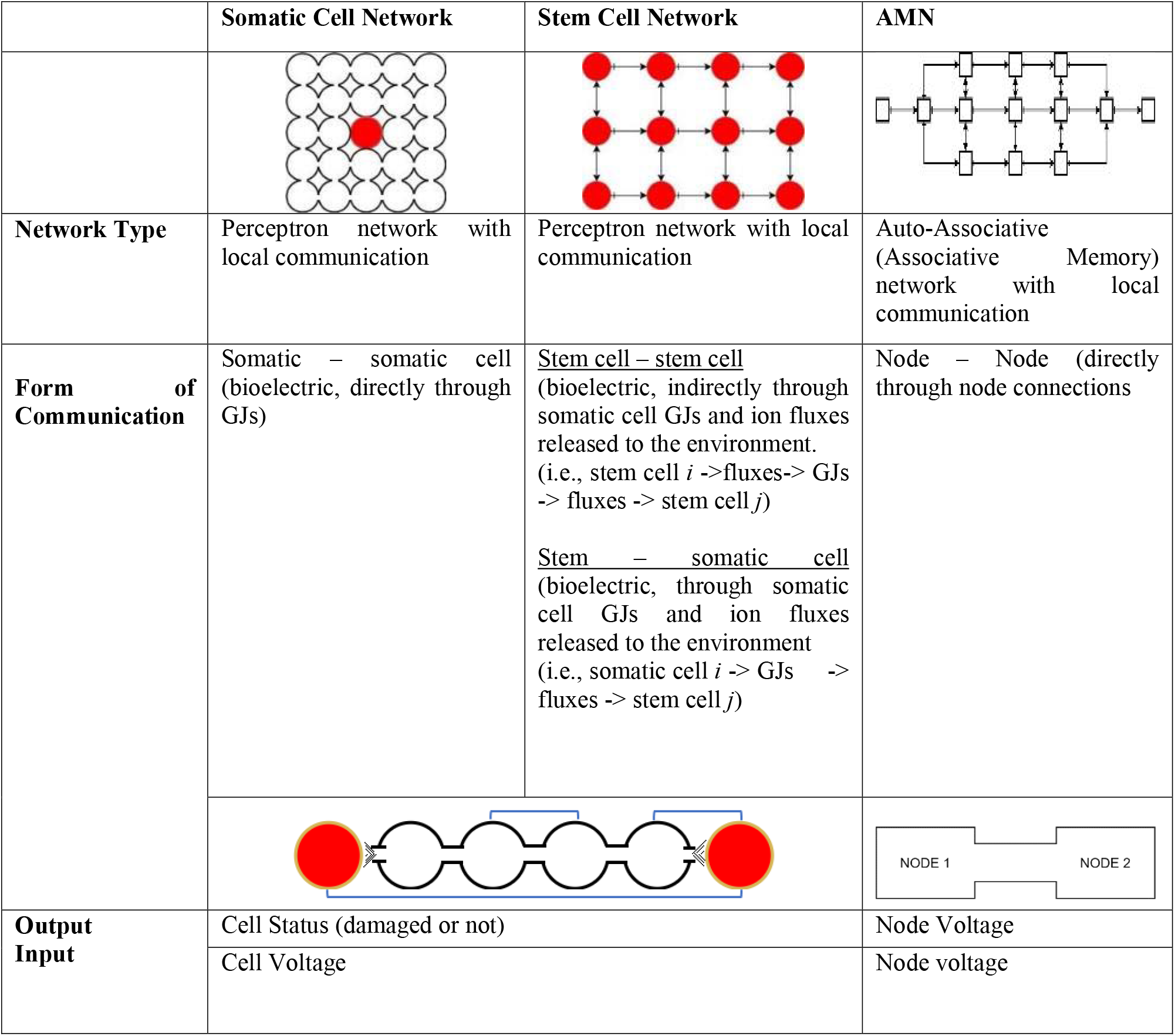
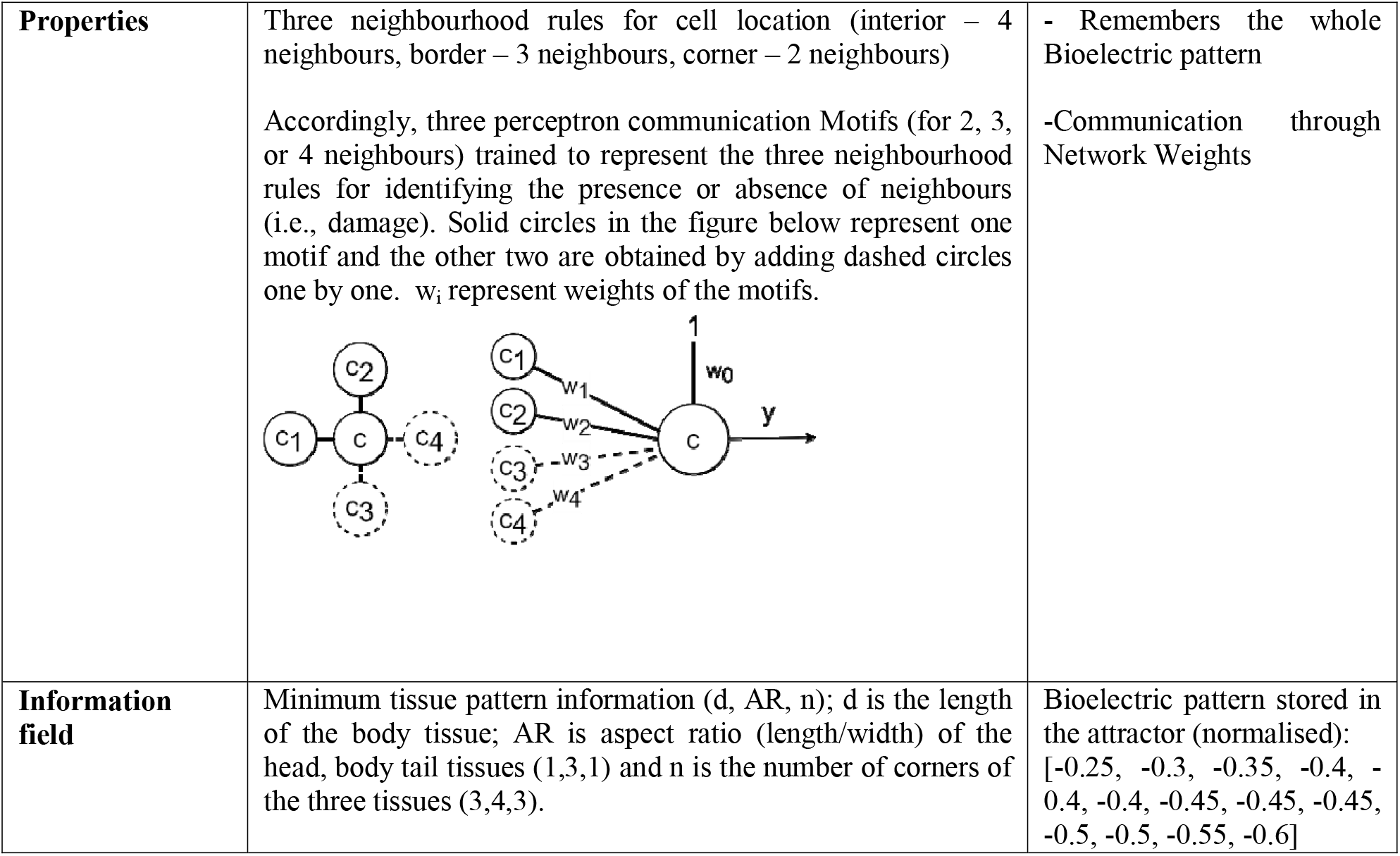
The three cell networks in the framework and their structure, properties and activities

|  | Somatic Cell Network | Stem Cell Network | AMN |
| --- | --- | --- | --- |
| Network Type | Perceptron network with local communication | Perceptron network with local communication | Auto-Associative (Associative Memory) network with local communication |
| Form of Communication | Somatic – somatic cell (bioelectric, directly through GJs) | <u>Stem cell – stem cell</u><br>(bioelectric, indirectly through somatic cell GJs and ion fluxes released to the environment. (i.e., stem cell $i \rightarrow \text{fluxes} \rightarrow \text{GJs} \rightarrow \text{fluxes} \rightarrow \text{stem cell } j$)<br><u>Stem – somatic cell</u><br>(bioelectric, through somatic cell GJs and ion fluxes released to the environment (i.e., somatic cell $i \rightarrow \text{GJs} \rightarrow \text{fluxes} \rightarrow \text{stem cell } j$) | Node – Node (directly through node connections |
| Output Input | Cell Status (damaged or not) |  | Node Voltage |
|  | Cell Voltage |  | Node voltage |

|  |  |  |
| --- | --- | --- |
| **Properties** | Three neighbourhood rules for cell location (interior – 4 neighbours, border – 3 neighbours, corner – 2 neighbours)<br>Accordingly, three perceptron communication Motifs (for 2, 3, or 4 neighbours) trained to represent the three neighbourhood rules for identifying the presence or absence of neighbours (i.e., damage). Solid circles in the figure below represent one motif and the other two are obtained by adding dashed circles one by one. $w_i$ represent weights of the motifs. | - Remembers the whole Bioelectric pattern<br>-Communication through Network Weights |
| **Information field** | Minimum tissue pattern information (d, AR, n); d is the length of the body tissue; AR is aspect ratio (length/width) of the head, body tail tissues (1,3,1) and n is the number of corners of the three tissues (3,4,3). | Bioelectric pattern stored in the attractor (normalised):<br>[-0.25, -0.3, -0.35, -0.4, -0.4, -0.4, -0.45, -0.45, -0.45, -0.5, -0.5, -0.55, -0.6] |

#### S1.2 Somatic cell network, communications and repair of local tissue damage

Somatic cells in each tissue form a somatic cell network. There are head, body and tail somatic cell networks consisting of perceptron neurons that communicate locally. For simplicity of computation, three formats or motifs are defined for this communication as shown in Fig.S1.

**Fig. S1.**
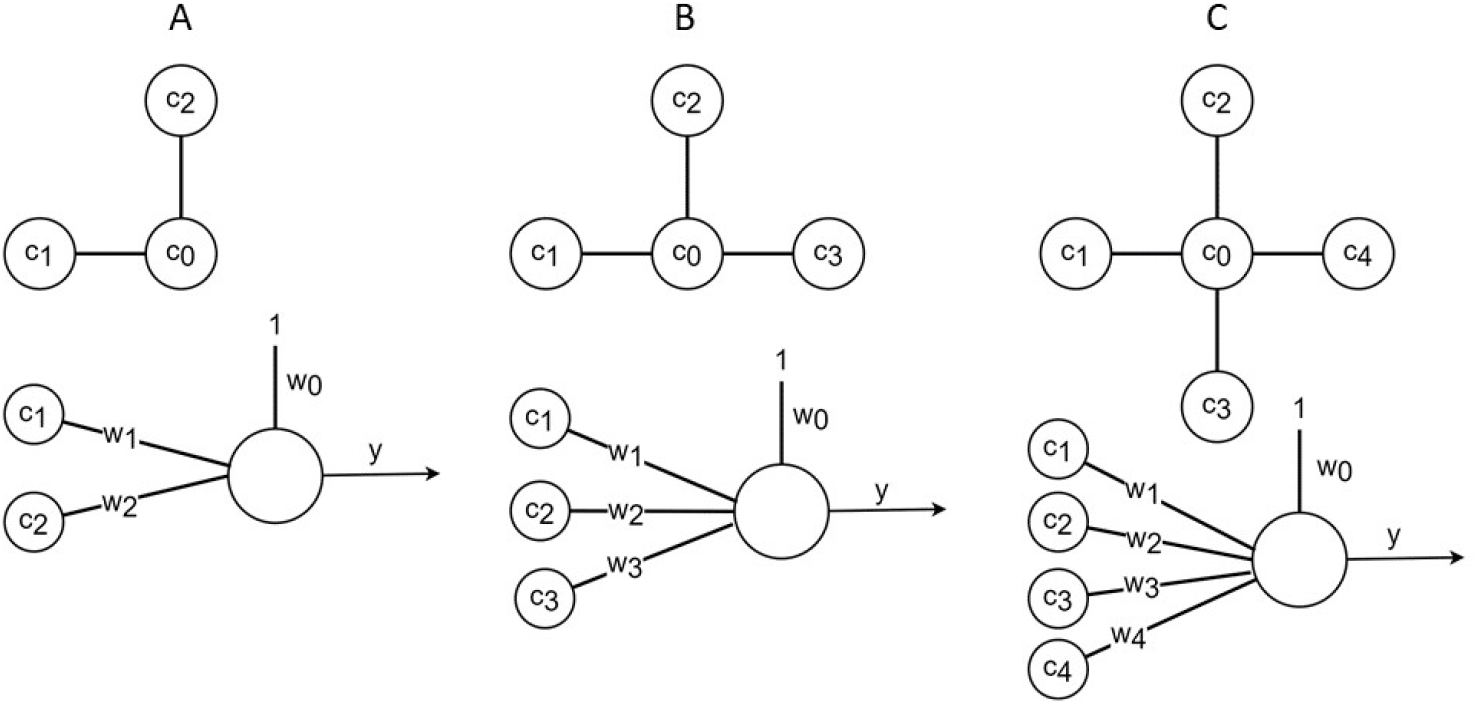
Perceptron motifs. Three communication motifs (A, B and C) depending on the number of neighbours 2, 3 or 4, respectively, in the somatic cell network. These denote corner (A), border (B) and interior (C) somatic cells. w_0_ denotes bias weight and w_i_ denotes weights associated with inputs.

##### The three perceptron communication motifs are

**Motif 1**: *<u>Rules for a Corner Cell</u>*: Two inputs [c_1_, c_2_] are normal indicative of no damage when [c_1_, c_2_] are non-zero negative (hyperpolarised) voltage values, and other input combinations indicate damage.

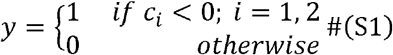

**Motif 2**: *<u>Rules for a Border Cell</u>*: Three inputs [c_1_, c_2_, c_3_] are normal when [c_1_, c_2_, c_3_] are non-zero negative voltage values, and other input combinations indicate damage.

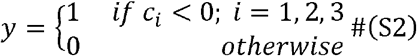

**Motif 3**: *<u>Rules for an Interior Cell</u>*: Four inputs [c_1_, c_2_, c_3_, c_4_] are normal when [c_1_, c_2_, c_3_, c_4_] are non-zero negative voltage values, and other input combinations indicate damage.

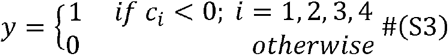

When there is local tissue damage with only missing somatic cells, the neighbours of the missing cells experience increased membrane voltage that triggers them identify the missing neighbours by applying the communication motif relevant to them (interior, corner or border cells). These cells with increased voltage become border of the damage. The stem cell(s) nearest to the damage also receive bioelectric communication from the affected somatic cells. This activates the stem cell network and upon finding that there are no missing stem cells, the nearest stem cell migrates to the damage border in the affected tissue and produces new somatic cells to restore the anatomy.

#### S1.3 Stem cell network, communication and repair of local and large scale damage

Stem cell network is an organism-wide network of stem cells represented by perceptrons with local communications. The same three communication motifs of the somatic cell networks apply to this network as well.

With respect to local stem cell damage, refer again to the example damage shown by the black square in Fig. 5A in the main text depicting one missing stem cell (and the surrounding tissue fragment). In this type of damage involving one (or more) stem cell, there still remain stem cells to the left, right, above and below the damage, indicative of local (in this case encased) stem cell damage. To identify all damage scenarios, we identify three generic patterns (primitives) that define the stem cell border of a damage as shown in Fig. 5B. The three primitives specifically refer to if there are stem cells: to the left, right or (above or below) the damage, respectively. Any encased damage would need all three generic pattern primitives and thus presence of all three primitives define the nature of all such local damages without whole tissue loss. For example, in the one stem cell damage case in Fig. 5A, all three patterns apply, as there are stem cells to the left, right and (above and below) the damage, indicating local (i.e., not whole tissue) damage. For other and large scale damages, not all three but one or two pattern primitives define the stem cell border. For example, Fig.5E shows stem cell border patterns (orange arrows) for whole tissue damages. Such damage is identified by the absence of one or more of the above three primitives in defining the stem cell damage border. The identification of damage patterns are specified below.

- ***Identification of local stem cell damage through primitives applied to the pattern of stem cell border (Figure 5B): All three primitives below should apply*** <u>Pattern 1</u>: stem cell damage border has one (or more) stem cells to the left of the damage in AP (along the body axis) direction. <u>Pattern 2:</u> stem cell damage border has stem cells above or below the damage (in DV direction). <u>Pattern 3:</u> stem cell border has one (or more) stem cells to the right of the damage (in AP direction).
- ***Identification of whole tissue damage through primitives applied to the pattern of stem cell border (Fig.5E): Only one of the above primitives should apply, not all three***.

When not all three pattern primitives described above are present in defining a damage border, it indicates whole tissue damage as in Fig.5E. For example, in the case of severed head tissue in Figure 5E, the stem cell damage border pattern has all stem cells on one side (left for the severed head tissue and right for the severed tail tissue) of the damage; therefore, only one primitive applies and 2 are absent indicating whole tissue damage.

We describe in some detail how these rules apply to identify local stem cell damage in the next section. Then we explain identification of large-scale stem cell damage involving loss of tissues in the following section.

##### Identification of local stem cell damage by the stem cell network

When the stem cell damage is local, the stem cell network finds the border of stem cell damage by applying perceptron motifs and the nature of local stem cell damage by applying the three pattern primitives described above. Specifically, for the damage case shown in Fig. 5A, the neighbours of the missing stem cell in the stem cell network sense a voltage increase beyond the 10% threshold, and by applying perceptron motifs the stem cell network identifies the four neighbours (circled in blue). These are the stem cells that have recognised that their neighbour stem cell is missing from the changes in their voltage status sensed through GJs of the intermediate tissue cells. These then form the border of stem cell damage. Then the above pattern primitives are applied to the damage border and it is evident that all three rules apply as there are border stem cells to the left, above and below, and right of the damage. This indicates local stem cell damage inside the stem cell network. Then the stem cell network regenerates a new stem cell to replace the missing one. Then, repair of damage to the tissue is accomplished in collaboration with the somatic cell network of the affected tissues that identify the damage border in the tissue by applying the perceptron motifs. The somatic cell damage border in the example case of local stem cell damage in Fig.5B is indicated by the yellow somatic cell border surrounding the damaged region (black square).

##### Identification of large-scale stem cell damage involving loss of whole tissues by the stem cell network

When the damage involves whole tissues, the stem cell network identifies the damage border and recognises that there is only one border pattern - all stem cells are on either side of the damage (Fig. 5E). Specifically, the three border pattern primitives are applied to the damage with the result that not all three are valid for this damage. From this, the stem cell networks recognises that a whole tissue(s) is missing. Now the question is how to identify which tissue is missing, head, body or tail, upon which stem cell network regenerates missing tissues while tapping into the information field for the minimum pattern information of lost tissues.

For identification of the type of missing tissues, we define few simple rules based on the communication between stem cells in the stem cell network with activation (positive) and inhibition (negative) signals as shown in Fig.2B. In this network, signals from tail to head (A/P direction) are negative (inhibitory) and in all other directions signals are positive (activation). We use these two features to identify the type of missing tissue(s) as follows:

###### Two rules of communication flow for identifying the type of missing whole tissues

- Stem cells in the damage border receive **positive signals** but **no negative signals**. This means there is no communication in the tail to head direction (left to right or P/A direction). Therefore,
  - If the border stem cells are in the head, then body is missing.
  - If the border stem cells are in the body, then tail is missing.
- Stem cells in the border receive **negative signals** but **no positive signals**. This means there is no communication from head to tail (right to left or A/P direction)
  - If the stem cells are in the body, then head is missing.
  - If the stem cells are in the tail, then body is missing.

##### Examples of application of the three stem cell border pattern primitives and two rules of communication flow for identifying global damages by the stem cell network

Below we illustrate in summary form how the stem cell network applies the three generic stem cell border pattern primitives and the above two rules of communication flow for identification of the type of missing whole tissues to recognise global damages.

<u>Example 1:</u> Whole tissue loss (*body and tail missing*): The head tissue separates after damage as shown in Fig. 5E (left) leaving body and tail intact. First, the stem cell network identifies the border of stem cell damage in the head tissue, which contains all stem cells on the head side. The network recognises that, of the three pattern primitives, only one applies. This indicates large scale tissue damage. Then, it identifies the type of missing tissue by applying the above rules of communication flow. As these border stem cells are in the head tissue, they **receive positive signals and no negative signals**. From this, the network identifies that the **body tissue is missing**. The stem cell network remaining in the head tissue regenerates body and tail by tapping into the information field for minimum tissue pattern information, upon which they produces body and tail stem cells which produce respective somatic cells to re-establish the stem cell network and somatic cell networks. Similarly, the body-tail part regenerates the head tissue and produces another worm.

<u>Example 2:</u> Whole tissue loss (*head and tail missing*): Let’s consider the body tissue remaining after damage in Fig.5E (middle) leaving three fragments. There are two damage regions: head side and tail side. For each region, the stem cell networks identifies the damaged stem cell border and recognises that there is only one stem cell pattern on either side. As not all three pattern primitives apply, the damage is due to tissue loss. Then, by applying the rules of communication flow to the stem cell border pattern in each damage region, the type of missing tissues are identified. For example, for the severed body tissue, the network identifies that one stem cell pattern **receives negative signals but not positive ones** (from head side) and vice versa for the tail side. Thus it identifies **head missing on the left side and tail missing on right side of damage**. The border stem cells then extracts the minimal pattern information for head and tail from the information field and the stem cells on the head side produce new stem cells which produce new somatic cells to regenerate the head using the minimal pattern information. Similarly, the stem cell network on the tail side produces new stem cells that regenerate the tail to the required pattern. Similar logic applies to the regeneration of the severed head and tail tissues. Single stem cell border pattern in these tissues defines whole tissue damage and absence of signals from the body side for the head fragment indicates missing body and tail and vice versa for the tail fragment. In both these cases, the respective border stem cells extract the minimal tissue plans from the information field and regenerate new stem cells to produce new somatic cells to match the pattern requirements for the missing tissues. Thus stem cell network fragments remaining after damage regenerate three worms from the three split segments.

#### S1.4 Associative Memory Network (AMN)

##### a. AMN training

The original HNN (Hopfield Neural Network) is a fully connected and symmetric (symmetric means weights for forward and backward interactions between two nodes are the same) recurrent network with two discrete values for states (*s*_*i*_ = ±1) When trained with Hebbian learning, these networks converge to at least one attractor. Our voltage input pattern vectors, however, are not discrete values of ±1 but a number of real (continuous) values. In [28], the authors used continuous state HNN with *s*_*i*_*∈* [−1,+1] and a fully connected asymmetric connection matrix. This continuous-state asymmetric network also trained well with Hebbian learning and converged into attractors from any inputs. Therefore, in our model, we use real-valued inputs, and we further explore beneficial and potentially meaningful connection configurations as an alternative to fully connected networks to reduce the computational burden of the network in having to work with a large number of weights.

Inputs in our model are real voltage values corresponding to the states of the 13 nodes in the network and scaled to *s*_*i*_ *∈* [−1,0] (minus values represent a hyperpolarised (more negative potential inside the cell than outside) voltage state of the cell membrane). Fig.8A (in the text) shows the original homeostasis bioelectric pattern in planaria represented by the 13 AMN nodes. The desired pattern of bioelectric gradient is in the range from −25mV to −60mV (milli Volts) which has been reported as the pattern of bioelectric homeostasis in planaria [29, 30]. In our model, the desired homeostasis bioelectric pattern for the organism consists of the 13 scaled voltage values of (−0.25, −0.3, −0.35, −0.4, −0.4, −0.4, −0.45, −0.45, −0.45, −0.5, −0.5, −0.55, −0.6) shown in Fig.8A. These correspond to the voltage values of the 13 macro-level nodes and represent the desired network attractor. We trained the AMN with a large number of voltage patterns (approximate 600) randomly perturbed from the original voltage pattern in Fig.8A. On a side note, these perturbed voltage patterns can correspond to voltage patterns that can result from the perturbation of parameters in a set of Differential Equations (Eq. S4) representing bioelectric communication between somatic cells via gap junctions as in [31].

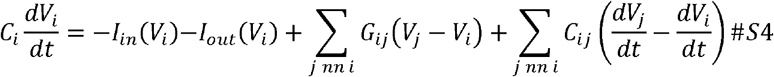

Where *V*_*i*_ is the membrane voltage of cell *i,G*_*ij*_ is the conductance of GJ between cells i and j (*C*_*ij*_ is capacitance), nn is the number of nearest neighbours of cell *i, I*_*in (or out)*_ (*V*_*i*_) is the current-voltage curve, and *C*_*i*_ is a constant; for details of these attributes refer to [31].

We simply highlight this ODE model to point out that the AMN in functionality (not in structure) represents existing bioelectric models of cell networks on a large scale (organism-wide). In future, this connection between the ANN and ODE models can be exploited to integrate cellular bioelectric and even molecular models into the framework (i.e., not only bioelectric models but also models of slow diffusion of molecules carrying signals for cell division in regeneration).

In training, the AMN starts with random connection weights that allow for learning the desired bioelectric pattern. Basically, we wish the AMN to store the target homeostasis bioelectric pattern in its attractor and invoke this pattern in response to any (altered) input voltage pattern vector. Therefore, all input patterns generated by perturbing the original voltage pattern should form the basin of attraction of the attractor. The learning algorithm goes through the following steps: (1) assignment of the initial states to the neurons (from the generated input patterns), (2) convergence of the network for a certain period (network simulation) followed by (3) application of Hebbian learning that adjusts weights incrementally until the network learns (stores) the target voltage pattern in an attractor and correctly recalls it from any input voltage condition that the organism may encounter.

The model dynamics use asynchronous neuron state updates with a rectilinear function (ReLU) in the nodes (Fig.S2) that calculates the output (voltage) *s*_i_ of *i*^th^ node as follows (Eq. S5):

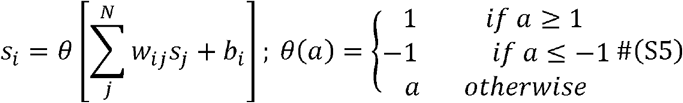

where *w*_*ij*_ is the connection weight between node *i* and *j, θ* is the transfer function (ReLU) and *b*_*i*_ is the bias weight of node *i*. In order to be able to store the desired pattern, we set random values for the connection weights (*w, b*) and adjusted them using Hebbian learning rule:

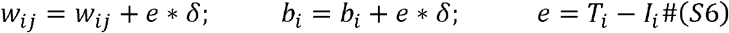

where *δ* is the learning rate, *e* is the difference between target pattern *T* and current input pattern *I*.

**Fig.S2.**
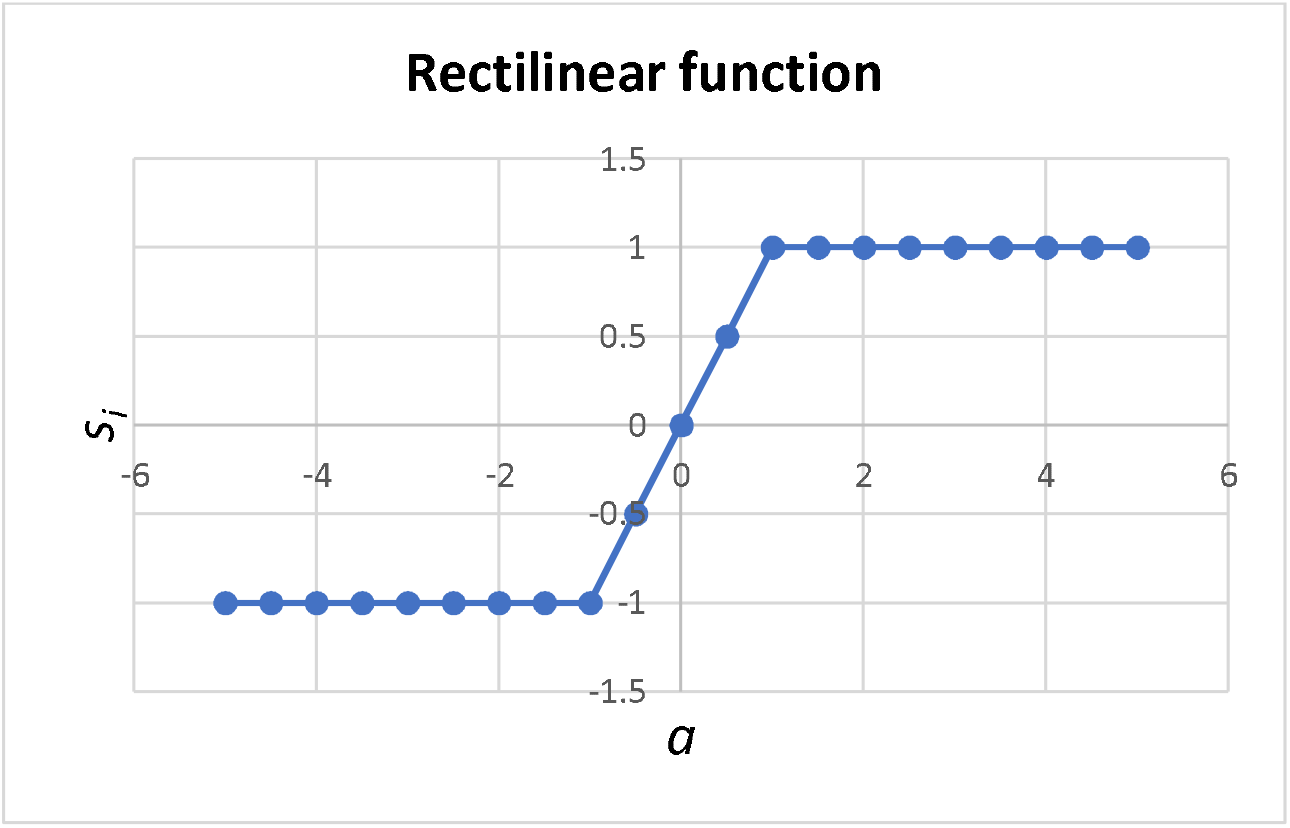
Rectilinear function for output (voltage) s_i_ of a node in AMN.

We designed the structure of the network by iteration. We first used a fully connected network as in HNN where all nodes communicate with each other. As this requires a massive amount of calculation to find an attractor with a large number of connection weights, we then reduced the connectivity in a number of stages to explore a simpler yet meaningful connection configuration that stores and recalls the pattern with minimum computation. We found a minimal connection configuration where local communication prevails as in Fig.S3 and where both the computation and memory storage requirements are drastically reduced. This resulted in 40 connection weights, a reduction of 87% from 312 connection weights in the fully connected network. This means that the reduced network has only 12% of the connections of the full network. The trained weights of the AMN are given in Table S3.

The minimal network topology that remembers and recalls the original homeostasis bioelectric pattern is as follows:

- Connections from head to tail and in the transverse (from the midline to edges) directions are positive (activation signals).
- Connections from tail to head are negative (inhibitory signals).

**Fig. S3.**
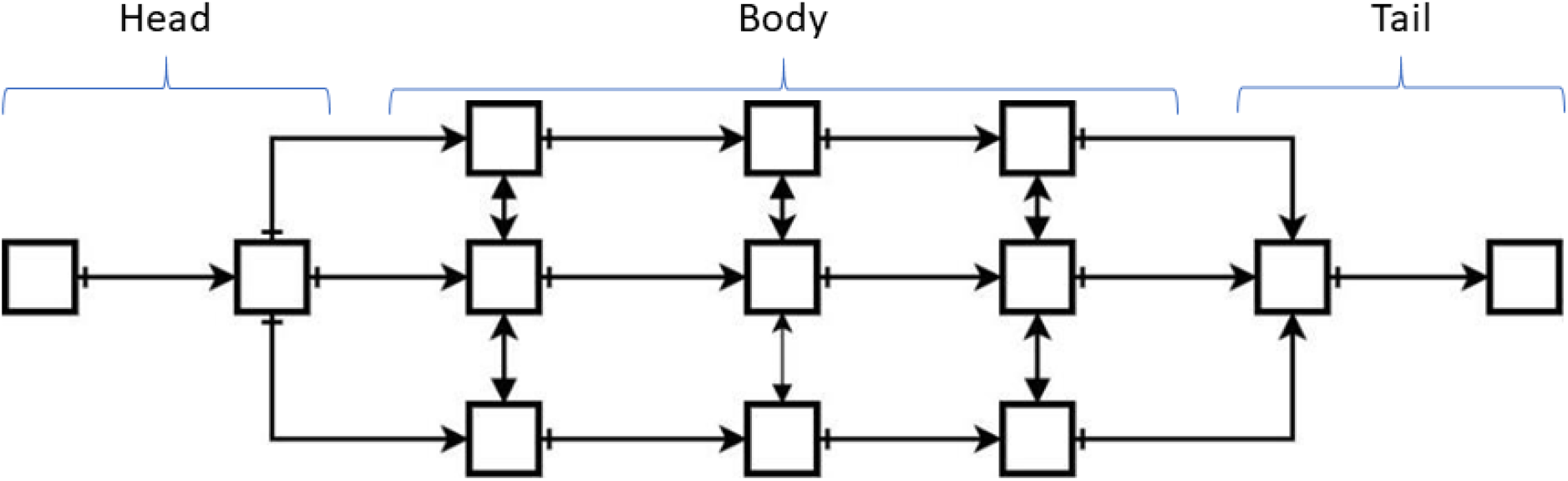
A minimal AMN structure that stores and recalls the original bioelectric pattern. Activation signals from head to tail (A/P) and transverse directions and inhibitory signals from tail to head (P/A) direction.

The fact that such a simplified and organised structure emerged was intriguing in itself. Specifically, this communication structure means that a node only receives signals from adjacent cells as follows:

- **For Anterior-posterior (AP) axis (longitudinal direction of the body):**
  - Positive signals: flow from head to tail
  - Negative signals: flow from tail to head
- **For the Dorsal -ventral (DV) axis (transverse direction of the body):**
  - Positive signals flow in both directions from mid-body to edge

Recall that this communication structure between nodes was adopted for the stem cell network as well as described in the text. Further, the above signal flow structure along and across the whole organism was used to derive the rules for identification of the type of missing whole tissues by the stem cell network as presented in the previous section.

##### b. AMN maintains bioelectric homeostasis under regular perturbations

In this section, we first discuss details of the algorithms and computations involved in the restoration of bioelectric homeostasis under normal physiological conditions and then present those involved in its restoration after recovery from damage.

The voltage of each AMN node ***V***_***k***_ is calculated by the following formula:

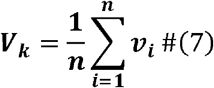

where n is the number cells in a node (e.g., n=300 cells), ***v***_***i***_ is the voltage of cell *i*, and *k* is the node number *k ∈* [1,13]. Fig. 3A (and 8A) in the text shows the unperturbed original bioelectric pattern. Fig.3B shows an example of perturbed voltage in a single node where one or few of its cells experience voltage change, and Fig.3C shows the perturbed voltage in all nodes where all cells experience voltage changes.

###### b (i) A single cell in a node receives perturbation to its voltage

Cells may be affected by their surroundings, such as temperature, light, chemicals and processes inside the cells, during normal physiological function, causing the voltage to change. According to the literature, membrane voltage can change up to ± 10% during normal operation [26]. We assume a threshold of ±10% for normal perturbations of membrane voltage of a cell.

Let ***∈*** be the voltage change in a single cell. The updated voltage in node k is presented as (e.g., Fig.3B):

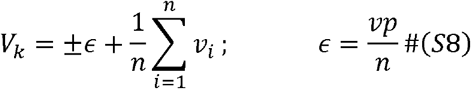

where, ***p*** is the percentage of voltage change, e.g. −10% to 10%, and v is the original voltage of a single cell.

###### b (ii) Many cells receive voltage perturbations

In the case where many cells undergo change in voltage (e.g., Fig.3C), the total voltage change in a node N_k_ (k=1, 2, .., 13) is computed as follows:

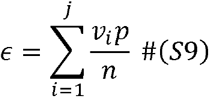

where j is the number of cells in node N_k_ receiving voltage perturbations

The algorithm for AMN recovery of the original voltage pattern from perturbed voltage vector is as follows:

- Determine the current altered voltage vector for the nodes (using Eq. S8 or S9).
- Present the current voltage pattern as input to AMN
- AMN iterates a number of times until it reaches a steady state which is the original bioelectric pattern stored in the attractor.
- In normal physiological functioning, the network is assumed to make affected nodes (specifically, affected cells within nodes) incrementally alter the bioelectric state until they return to the normal state. In the real organism, this could be achieved by the bioelectric activation of the required molecular signalling networks that manipulate membrane bound ion channels to adjust the cell voltage.

##### c. AMN maintains bioelectric homeostasis after regeneration

The AMN also restores the bioelectric pattern after the stem cell network completes regeneration. Recall that an AMN node consists of 300 somatic cells and 12 stem cells. In case of small damages where any single AMN node is not completely damaged (i.e., not all cells within a node get damaged), the AMN can restore bioelectric state after regeneration of missing somatic or stem cells in individual nodes. However, when an AMN network node gets damaged, the spatial topology of the network is broken. The network will then lose the node and all connection weights to this node. After completing regeneration, the connections from this node to others need to be re-established to restore lost connection weights. Further, the voltage of repaired nodes will not be the same as the required and therefore, nodal voltage also needs to be updated. We assume that after the initial training of the AMN, the original bioelectric pattern (bioelectric gradient) captured by the attractor is stored in the Information Field along with the connection topology as a template. After nodes are repaired following injury, the AMN (Fig.S3) retrains itself by accessing the bioelectric pattern from the field until the network learns to store it in the attractor and the connections and weights to repaired nodes are re-established. The algorithm is as follows:

- Initialise connections: Connections from the new node to others are re-established based on the network topology, and corresponding weights are initialised with random values (e.g., 0.1 for positive, −0.1 for negative signals).
- Retrain the AMN: The network retrains with the target bioelectric pattern accessed from the Information Field. Here the original pattern is perturbed to produce training patterns. (since the network has to train only the damaged connection weights, it is trained faster than the original network)
- Restore weights to repaired nodes.
- Adjust voltage: Cells in the affected node(s) alter the bioelectric state incrementally until they return to the normal equilibrium state.

In the text, we describe the operation of the whole framework integrating the three levels.

### S3. Some Results

**Table S2:** Weights of perceptron motifs

|  | **Bias** | **Input 1** | **Input 2** | **Input 3** | **Input 4** |
| --- | --- | --- | --- | --- | --- |
| **Motif 1** | -0.01 | 0.01 | 0.01 |  |  |
| **Motif 2** | -0.2 | 0.1 | 0.1 | 0.1 |  |
| **Motif 3** | -0.05 | 0.01 | 0.01 | 0.03 | 0.01 |

**Table S3:** Weights of Associative Memory Network (AMN)

| **Node** | **1** | **2** | **3** | **4** | **5** | **6** | **7** | **8** | **9** | **10** | **11** | **12** | **13** |
| --- | --- | --- | --- | --- | --- | --- | --- | --- | --- | --- | --- | --- | --- |
| **1** | 0.0 | 0.01 | 0.0 | 0.0 | 0.0 | 0.0 | 0.0 | 0.0 | 0.0 | 0.0 | 0.0 | 0.0 | 0.0 |
| **2** | -0.86 | 0.0 | 0.01 | 0.01 | 0.01 | 0.0 | 0.0 | 0.0 | 0.0 | 0.0 | 0.0 | 0.0 | 0.0 |
| **3** | 0.0 | -1.0 | 0.0 | 0.01 | 0.01 | 0.01 | 0.0 | 0.0 | 0.0 | 0.0 | 0.0 | 0.0 | 0.0 |
| **4** | 0.0 | -1.0 | 0.01 | 0.0 | 0.0 | 0.0 | 0.01 | 0.0 | 0.0 | 0.0 | 0.0 | 0.0 | 0.0 |
| **5** | 0.0 | -1.0 | 0.01 | 0.0 | 0.0 | 0.0 | 0.0 | 0.01 | 0.0 | 0.0 | 0.0 | 0.0 | 0.0 |
| **6** | 0.0 | 0.0 | -1.0 | 0.0 | 0.0 | 0.0 | 0.01 | 0.01 | 0.01 | 0.0 | 0.0 | 0.0 | 0.0 |
| **7** | 0.0 | 0.0 | 0.0 | -1.0 | 0.0 | 0.01 | 0.0 | 0.0 | 0.0 | 0.01 | 0.0 | 0.0 | 0.0 |
| **8** | 0.0 | 0.0 | 0.0 | 0.0 | -1.0 | 0.01 | 0.0 | 0.0 | 0.0 | 0.0 | 0.01 | 0.0 | 0.0 |
| **9** | 0.0 | 0.0 | 0.0 | 0.0 | 0.0 | -1.0 | 0.0 | 0.0 | 0.0 | 0.01 | 0.01 | 0.01 | 0.0 |
| **10** | 0.0 | 0.0 | 0.0 | 0.0 | 0.0 | 0.0 | -1.0 | 0.0 | 0.01 | 0.0 | 0.01 | 0.0 | 0.0 |
| **11** | 0.0 | 0.0 | 0.0 | 0.0 | 0.0 | 0.0 | 0.0 | -1.0 | 0.01 | 0.0 | 0.0 | 0.01 | 0.0 |
| **12** | 0.0 | 0.0 | 0.0 | 0.0 | 0.0 | 0.0 | 0.0 | 0.0 | -1.0 | -1.0 | -1.0 | 0.0 | 0.01 |
| **13** | 0.0 | 0.0 | 0.0 | 0.0 | 0.0 | 0.0 | 0.0 | 0.0 | 0.0 | 0.0 | 0.0 | -1.0 | 0.0 |
| **Bias** | -0.243 | -0.5 | -0.634 | -0.688 | -0.688 | -0.732 | -0.837 | -0.837 | -0.83 | -0.936 | -0.936 | -1.99 | -1.145 |

#### S3 Further examples of implementation of the regeneration framework – Recovery from complex damage cases

This section presents few more examples of small and large local damages and few complex and severe global damage cases and how the framework successfully and completely regenerates the whole worm.

##### Case 1: Multiple somatic cells missing

The case of more than one somatic cell missing is similar to one cell missing (Fig.S4). However, more than one stem cell may check for damage and implement recovery. Fig.S4A illustrates an example case where five somatic cells (black) have been lost. The somatic cell network determines the border of damage (yellow colour cells in Fig. S4B. Here, three nearest stem cells recognise that there constituent somatic cells have been damaged (Fig.S4B burgundy cells) and recognise what/where the damage is. Here, only somatic cells are gone, and each stem cell repairs the tissue in its neighbourhood (Fig.S4C), and then AMN restores the original voltages (Fig.S4D).

**Fig. S4.**
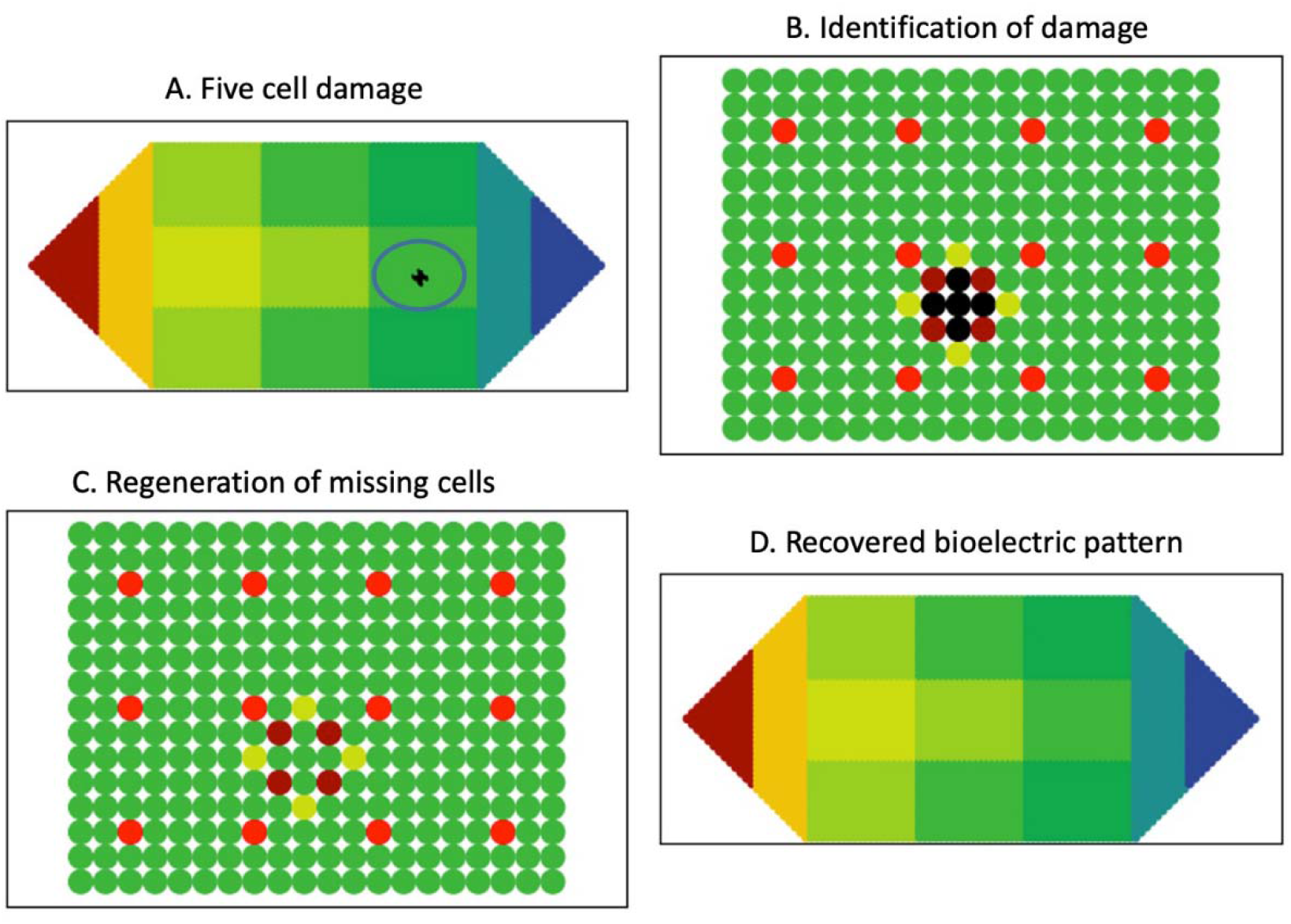
An example of five somatic cell damage – (A) Five damaged somatic cells (black cells); (B) Damage identification: damaged cells and their neighbours (yellow colour cells; yellow indicates that voltage in the region is higher than normal); burgundy cells and red cells respectively are stem cells in the affected area and unaffected stem cells; (C) Regeneration of missing cells; the voltage of new cells are equal to the voltage of the producing (burgundy) stem cells; (D) AMN recovers the original bioelectric state

##### Case 2: Organism recovers from damage to many stem cells and surrounding tissue – an AMN node damage

###### Damage identification and recovery

This case involves a relatively large damage (about 300 cells) involving 12 stem cells and surrounding somatic cells in the interior of the body as shown in Fig.S5A and in the magnified view of damage region (black square) in Fig.S5B. This is damage to a node in the AMN. The methods of sensing, identifying and repairing the damage are similar to the methods for the previously presented case of a stem cell and surrounding tissue damage. Specifically, the stem cell network first identifies missing stem cells and the damage border in the stem cell network (burgundy colour dots (stem cells) surrounding the black square in Fig.S5B). As all three pattern primitives apply, the stem cell damage is recognised as local. The somatic cell network identifies missing neighbour somatic cells and identifies the damage border in the tissue (yellow cells surrounding black square), using the perceptron motifs. Fig.S5C-D show few snapshots of regeneration for this damage case. Specifically, the border stem cells migrate to the damage site and produce 12 new stem cells that generate somatic cells to fill the space left by the damage.

###### Restoration of biolectric homeostasis

As a node in the AMN is gone, we apply the algorithm in section S1.4c above to restore bioelectric homeostasis.

**Fig.S5.**
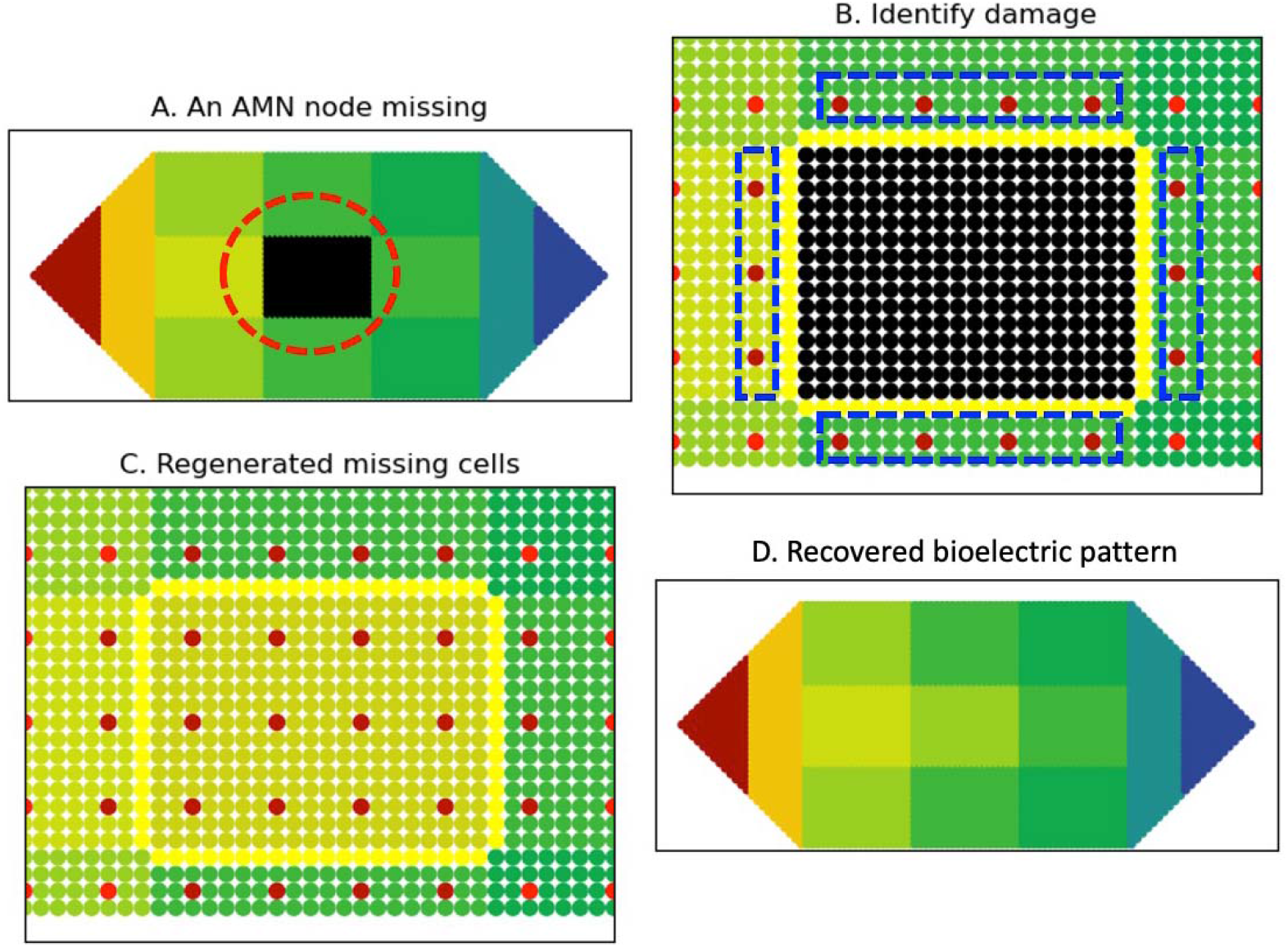
Large damage of the size of an AMN node including 12 stem cells and 300 somatic cells. (A) Damage region (black square) where cells are missing; Dash lines highlight the affected cell area; (B) Somatic cell network identifies border surrounding tissue damage (yellow border) and stem cell network identifies its damage border (burgundy dots surrounding the damage); yellow and burgundy indicate that voltage in the region is higher than normal); (C) damage area and surrounding tissue after regeneration (D) AMN restores the body-wide bioelectric pattern.

##### Case 3: Organism recovers from separated whole tissues along with interior damage

Fig.S6 depicts an injury severing head and tail from the body and further incurring damage to the interior of the body of the worm (Fig.S6A). This is a combination of three damages: head, tail, and a group of interior body cells (AMN node in this case) missing. Figure S6B shows the group of cells removed from the body of the worm.

For the damage in Fig.S6A, the three combined damages mean that it requires a combination of three corresponding recovery processes. Using the three border pattern primitives, stem cells identify the two ‘whole tissue’ damages and the ‘local damage’ to the tissue. The worm achieves head and tail regeneration as in the Case 3 – Head regenerates body and tail shown in the text and missing piece regeneration as in Case 2 (above) for missing AMN node. Fig.S7 shows few snapshots in the repair of the worm starting from the remaining body tissue as in Fig.S6A and the final step of restoration of the voltage pattern.

**Fig.S6.**
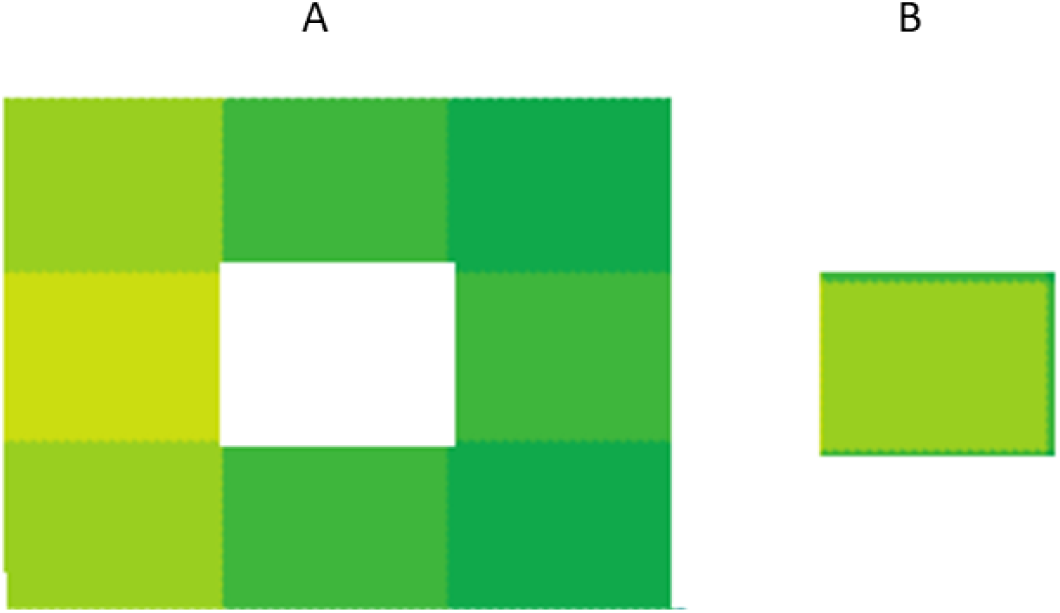
Complex damage: A) Separation of parts (head and tail) and a portion of the body interior; B) portion separated from the body interior (A block represents an AMN node)

Fig.S8 shows the regeneration of the full worm from the small interior part separated from the body as in Fig.S6B. The worm achieves this by applying the same procedures as in Case 3 – Head regenerates body and tail (in the text). Specifically, after determining the border and missing tissues (Stages 2, 3), the stem cells proceed to restore the head and tail (stage 4) (Fig.S8A). Head and tail regenerate concurrently starting from small tissues as a small worm (Figure S8B-D) that grows into the exact original form following shape information taken from the Information field (Fig.S8E). After regeneration, the AMN is re-established, and the bioelectric pattern is restored (Fig.S8F) (stage 5).

**Fig. S7.**
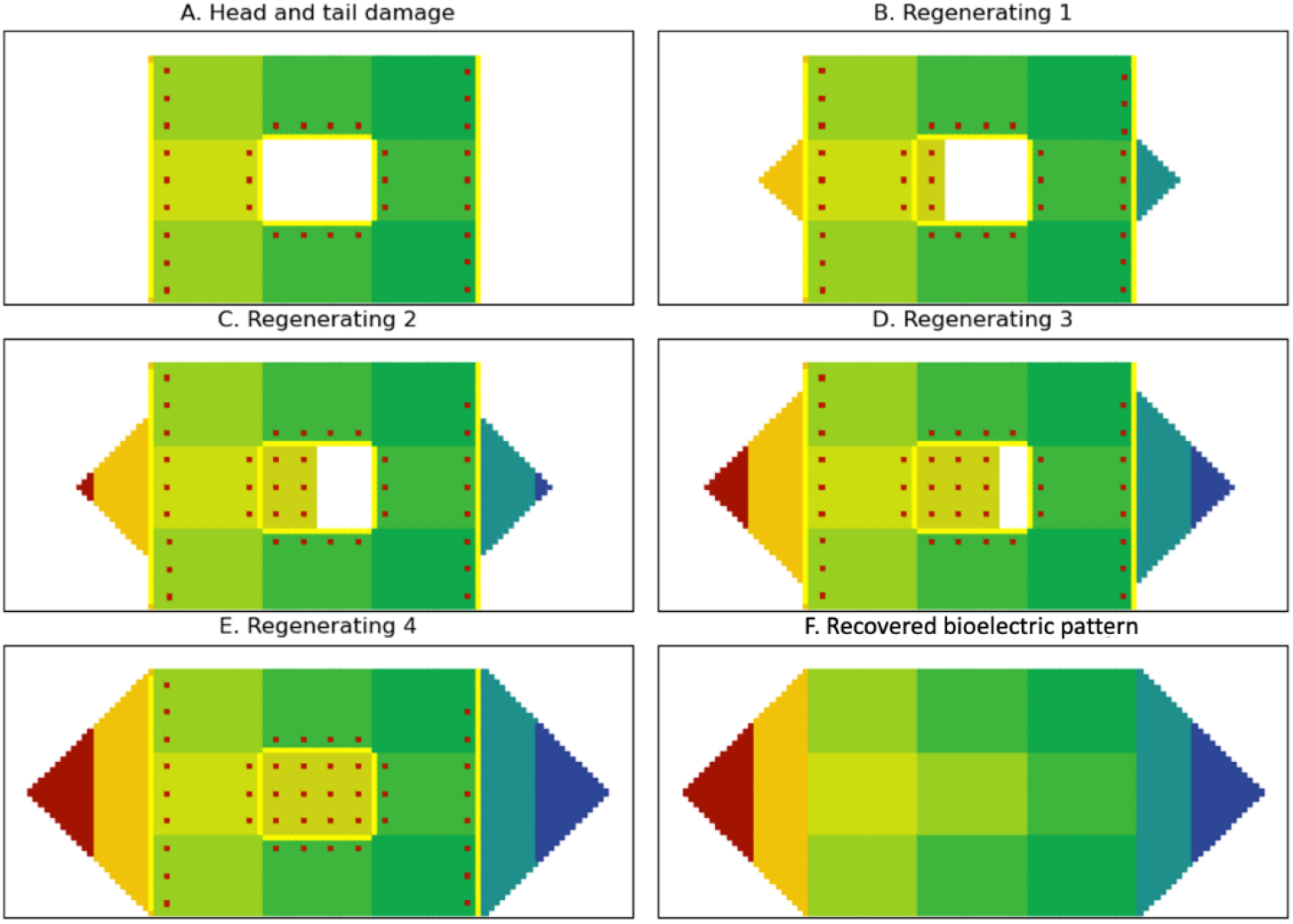
Few snapshots from the process of regeneration from damage in Fig.S6A.

**Fig.S8.**
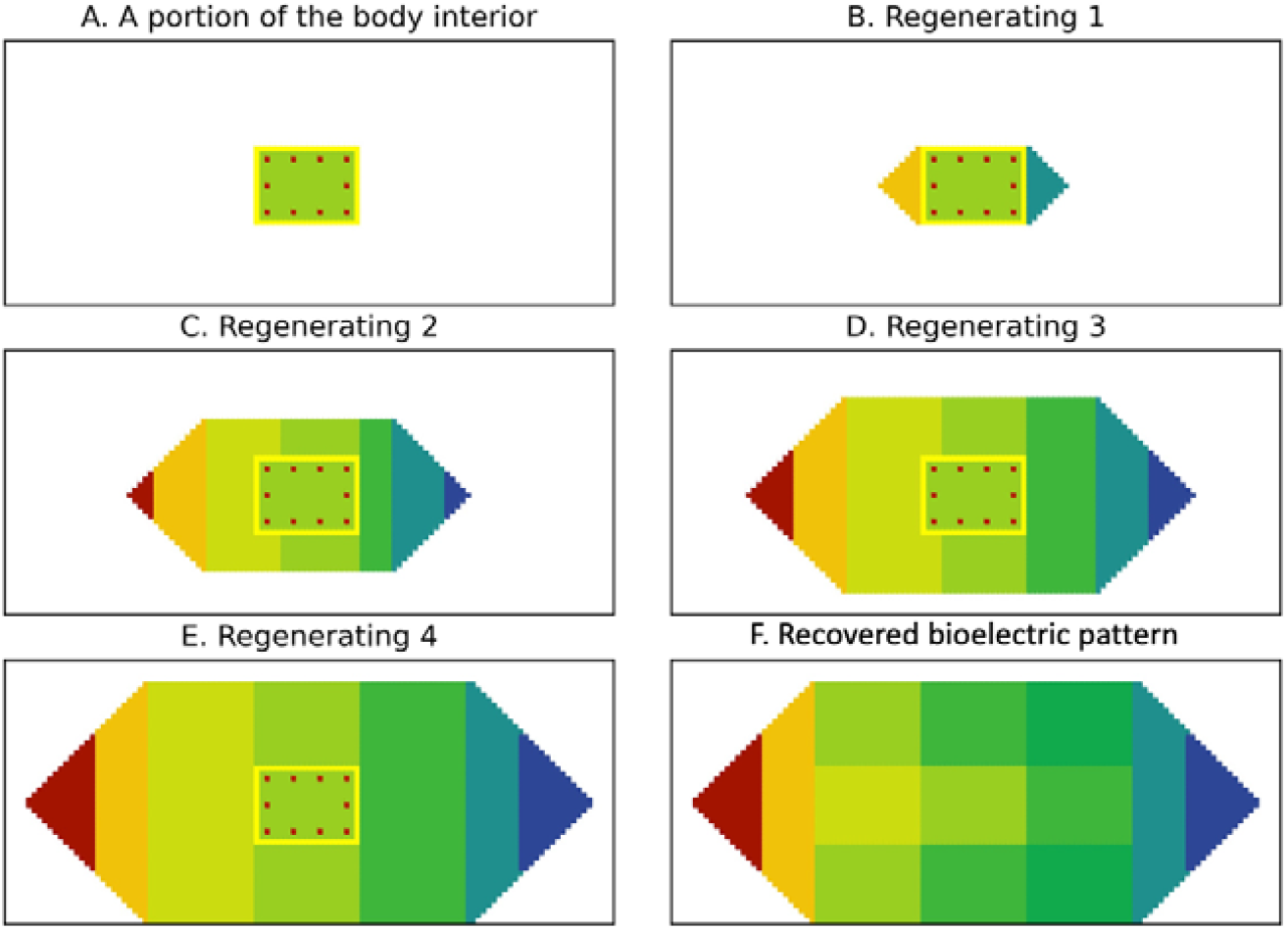
Regeneration of the full worm from the small fragmented part in Fig.S6B.

##### Case 4: Complex damage boundary

Fig.S9 presents another complex damage case where the worm is split into two with a vertical cut part-way through the body and then a horizontal cut through the body and tail: one part consists of complete head tissue and parts of body and tail tissues and the other part consists of the rest of the body and tail parts.

**Fig.S9.**
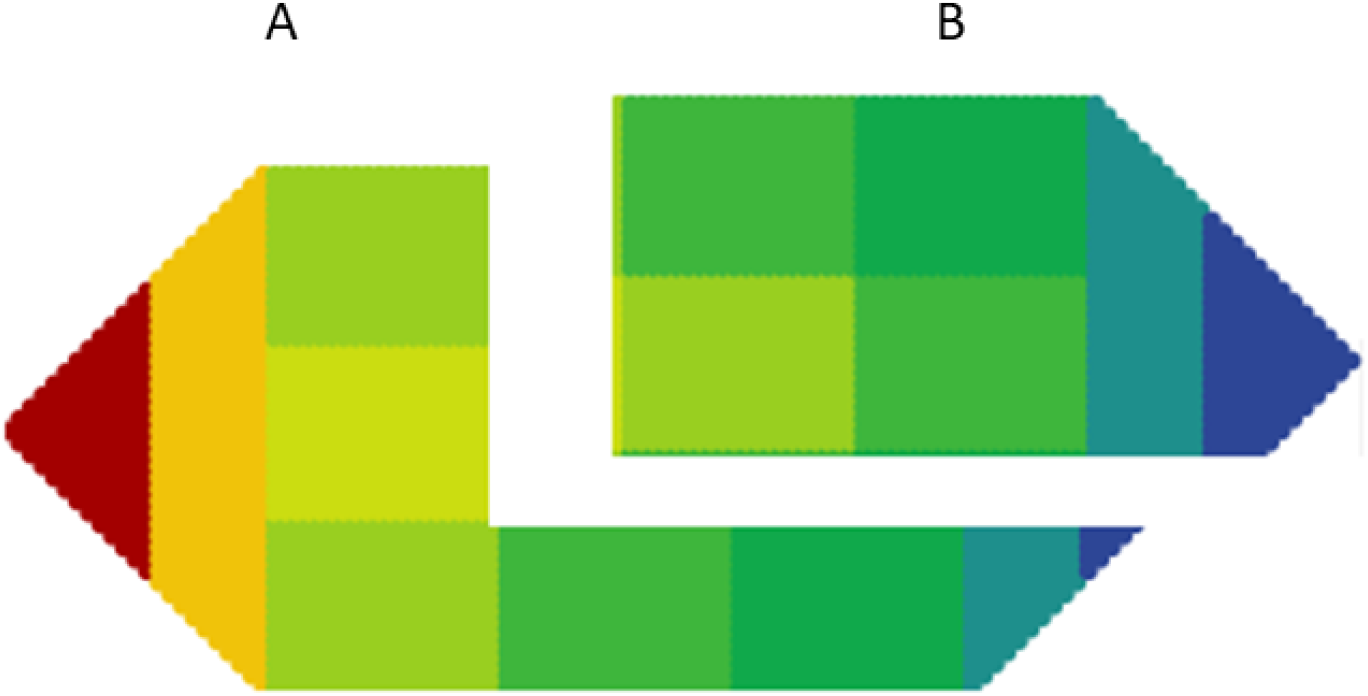
Complex damage splitting the worm into two through a number of issues. (A) Part 1: Intact head with body and tail damage. (B) Part 2: No head and body and tail damage.

###### Part 1

As shown in Fig.S9A, after damage, there is still the complete head, about half the body, and a portion of the tail tissue left. As such, this is not whole tissue damage. Stem cells identify and repair damage as the case of ‘many stem cells missing’. The shapes of the body and tail tissues are determined from the current borders and length of sides (d) accessed from the Information field. Regeneration steps for this case are shown in Fig. S10.

###### Part 2

There is still about half the body and a large portion of tail tissue left after damage (Fig.S9B). As such, this is a combination of whole and partial tissue damage. The process of damage detection and identification is as in the above cases. Specifically, stem cells identify the head missing (whole tissue loss) and damage to the body and tail parts (local/partial damage). Stem cells get pattern information for the new head from the information field (length (d), aspect ratio AR, corners (n)) and regenerates a head. The repair of the body and tail relies on the pattern information retained in the damaged tissue and extracting the length and width information from the field. Regeneration steps of this damage are shown in Fig.S11.

**Fig.S10.**
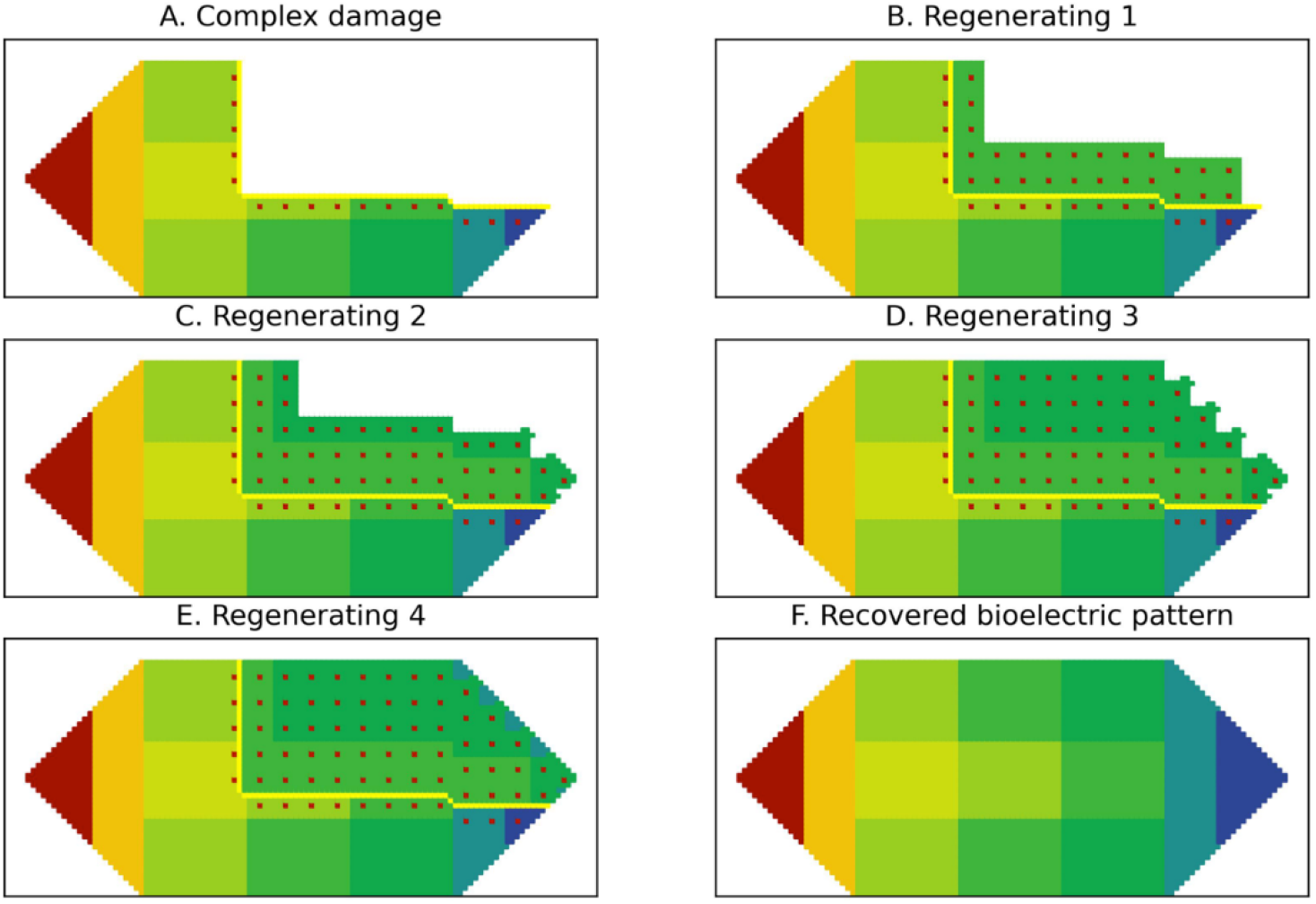
Regeneration process for damage in Fig.S9A.

**Fig.S11.**
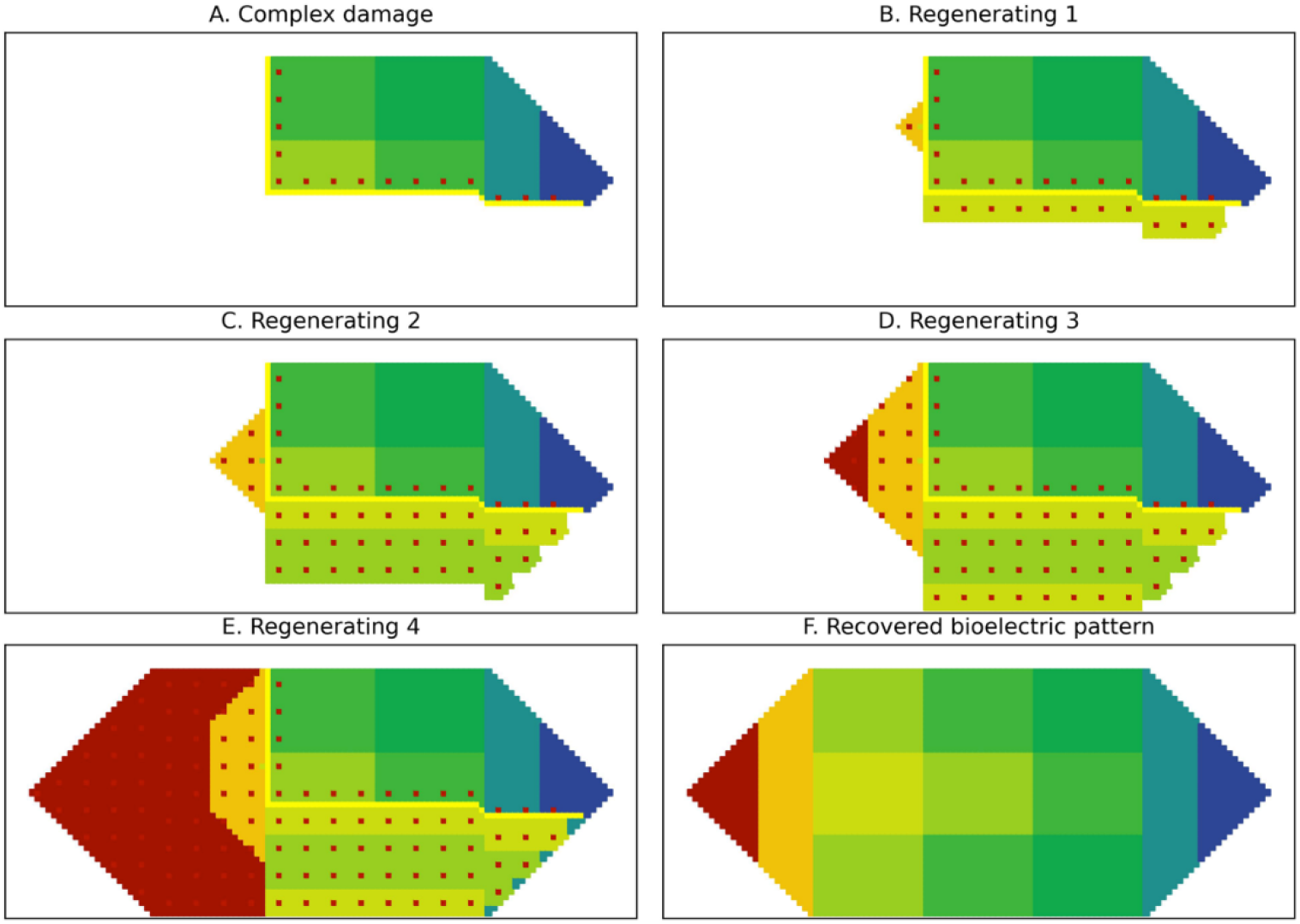
Regeneration steps for the damage in Fig.S9B.

